# Neurophysiological Correlates of Phase-Specific Enhancement of Motor Memory Consolidation via Slow-Wave Closed-Loop Targeted Memory Reactivation

**DOI:** 10.1101/2024.01.16.575884

**Authors:** Judith Nicolas, Bradley R. King, David Levesque, Latifa Lazzouni, David Wang, Nir Grossman, Stephan P. Swinnen, Julien Doyon, Julie Carrier, Geneviève Albouy

## Abstract

Memory consolidation can be enhanced during sleep using targeted memory reactivation (TMR) and closed-loop (CL) acoustic stimulation on the up-phase of slow oscillations (SOs). Here, we tested whether applying TMR at specific phases of the SOs (up vs. down vs. no reactivation) could influence the behavioral and neural correlates of motor memory consolidation in healthy young adults. Results showed that up- (as compared to down-) state cueing resulted in greater performance improvement. Sleep electrophysiological data indicated that up-stimulated SOs exhibited higher amplitude and greater peak-nested sigma power. Task-related functional magnetic resonance images revealed that up-state cueing strengthened activity in - and segregation of - striato-motor and hippocampal networks; and that these modulations were related to the beneficial effect of TMR on sleep features and performance. Overall, these findings highlight the potential of CL-TMR to induce phase-specific modulations of motor performance, sleep oscillations and brain responses during motor memory consolidation.

## 1. Main

Memory consolidation is the process occurring offline, between practice sessions, by which labile memory traces become more robust ^1^. Seminal rodent work suggests that consolidation relies on the strengthening of the mnemonic representations by the spontaneous reoccurrence - during post-learning offline periods - of hippocampal firing patterns associated with the initial encoding ^2–4^. This reactivation process has been particularly studied during post-learning sleep and there is consistent evidence that Non-Rapid Eye Movement sleep (NREM) oscillations such as slow oscillations (SO - high amplitude oscillations in the 0.5–2 Hz frequency band) and spindles (short burst of oscillatory activity in the 12–16 Hz sigma band) orchestrate the spontaneous occurrence of these hippocampal reactivations ^1,5^. Spontaneous reactivations of task-related brain patterns have since been observed during post-learning sleep in humans after both declarative and motor learning (see ^6^ for a review). The field has recently seen a surge of research examining whether experimental interventions can induce these reactivations in the human brain and eventually enhance the memory consolidation process ^7,8^.

An experimental intervention that has shown promise to enhance memory consolidation is Targeted Memory Reactivation (TMR) ^9^. TMR is a non-invasive procedure which consists of replaying, offline, sensory stimuli that were previously associated to the task during initial memory encoding ^9,10^ Auditory TMR applied during post-learning NREM sleep has been consistently shown to boost both declarative and motor memory consolidation in healthy young adults ^e.g., 11–15^ and this process is thought to be mediated by the modulation of SO ^14^ and spindle ^12,16^ characteristics as well as their coupling ^14^. Inspired by studies showing that auditory clicks delivered in a closed-loop (CL) fashion at the up-state of the SO can optimize declarative memory consolidation ^e.g., 17^, recent studies have applied TMR at different phases of the SO (e.g., up- vs. down-stimulation) in an attempt to further optimize consolidation. Such CL-TMR interventions have been shown to increase SO and sigma band power following cues presented at the up- as compared to the down-phase of the SO ^18,19^ or as compared to a control night without stimulation ^20^. These studies show an overall memory advantage following up-state TMR ^18–20^, albeit performance does not always differ from all other stimulation conditions (e.g., from down- ^18^ or no-stimulation ^19^). Altogether, studies causally liking the specific SO phase of the reactivation to memory consolidation are sparse in the declarative memory domain and are non-existent in the motor memory domain. Additionally, the effect of slow-oscillation CL-TMR on the neurophysiological processes underlying memory reactivation and memory retention are poorly understood in both memory domains.

In this pre-registered study (https://osf.io/dpu6z)^1^, we used functional magnetic resonance imaging (fMRI) during task practice and electro-encephalography (EEG) during post-learning sleep to address these knowledge gaps and provide a comprehensive characterization of the neurophysiological processes supporting the effect of slow-oscillation CL-TMR on motor memory consolidation. Briefly, in a within-subject design, 31 young healthy participants learned 3 different motor sequences that were each associated to one specific sound during learning while their brain activity was recorded with fMRI (Pre-night, Figure 1a). During the subsequent post-learning night of sleep that was monitored with EEG (Night, Figure 1a), SOs were detected in real-time during NREM sleep and auditory cues that were associated to the motor learning task were delivered to specific phases of the SO reflecting either high or low brain excitability. Specifically, one sound was played at the peak of the SO (up-reactivated condition), another sound was played at the trough of the SO (down-reactivated condition), while the last sound was not replayed (not-reactivated, control, condition). To assess consolidation, motor task performance was retested on the three different conditions in the fMRI scanner the next morning (Post-night, Figure 1a). Our main results confirmed the pre-registered hypotheses as consolidation ^18^, SO amplitude ^17,19,21^, sigma band power ^17,19^ and task-related brain responses in hippocampo- and striato-cortical networks ^12,22^ were specifically boosted by SO-up-phase TMR.

**Figure 1:**
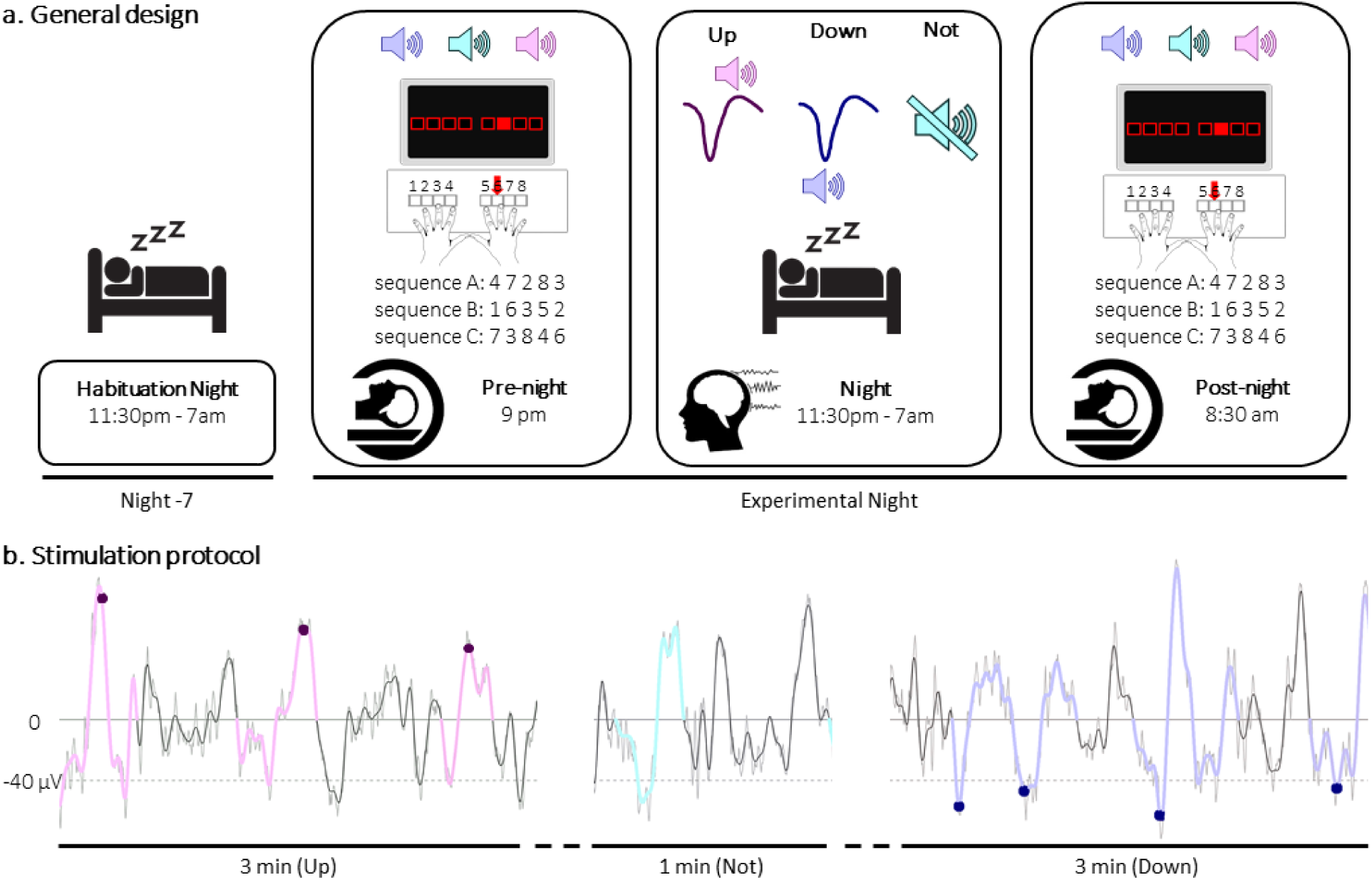
Experimental protocol. a. General design. Following a habituation night that was completed approximately one week prior to the experiment, 31 participants underwent a pre-night motor task session in the scanner, a full night of sleep in the sleep lab monitored with polysomnography during which slow-oscillation closed-loop targeted memory reactivation (CL-TMR) was applied, and a post-night retest session in the scanner. During the motor task (pre-night and post-night sessions), three movement sequences were performed (sequences A, B and C whereby 1 and 8 correspond to the right and left little fingers, respectively) and were cued by three different 100-ms auditory tones. For each movement sequence, the respective auditory tone was presented prior to each sequence execution. Two of these sounds were replayed during the post-learning sleep episode at specific phases of SO (up vs. down, see panel B for details) while the third sound was a control condition which was not replayed during the night. Note that the sequence / sound / condition combinations were randomized across individuals (see methods). b. Stimulation protocol. Sleep was recorded with EEG all night but recordings were monitored online for stimulation purposes during the first three hours of the night. The online SO detection algorithm was launched whenever the participant reached NREM2-3 stage. Three-min long up- and down-stimulation intervals alternated and were separated by 1-min no-stimulation intervals. The sounds associated to the up(or down)-reactivated sequence were then played on the peak(or trough) of the SOs within these alternating blocks. The algorithm performed the online detection on FPz. For down detection, a fast-moving average filter was employed with a window of 50 samples and a trough was detected when the signal went below a specific threshold adapted for biological sex ^23^ of −41µV in females and −39.5µV in males. For up-detection, the peak of a SO was identified when, in addition to the criterion described for trough detection above, peak-to-peak signal amplitude reached 77 µV in females and 74 µV in males (See methods for details). During up-stimulation intervals, the up-reactivated sequence sound (magenta dots) was played at the peak of each detected SO (up-stimulated SO / up-reactivated sequence) and during down-stimulation intervals, the down-reactivated sequence sound (blue dots) was played at the trough of each detected SO (down-stimulated SO / down-reactivated sequence). The third sound was not presented during the post-learning night (not-reactivated sequence). The colored oscillations in each panel represent the results of the offline SO detection algorithm that was used to compute, a posteriori, the accuracy of the online detection procedure and to detect SOs during rest intervals for further analyses (see methods). The online detection algorithm was manually stopped when the experimenter detected REM sleep, NREM1 or wakefulness and thus no stimulation was sent.

## 2. Results

### 2.1. The effect of TMR on motor performance depends on the phase of the stimulated SO

We tested whether the stimulation conditions (up-, down- and not-reactivated) influenced the behavioral index of motor memory consolidation, i.e., the offline changes in performance observed between the pre-night test session and the beginning of the post-night training session (see Figure 2a). Results show that offline changes in performance differed depending on the phase of the stimulated SO (Condition effect (F (2,54) = 3.9, p = 0.027 (0.034 sphericity corrected), η² = 0.13; Figure 2b, n = 28). Specifically, offline changes in performance were greater for both the up- and not-reactivated sequences as compared to the down-reactivated sequence (up vs. down: t = 2.32, p-value = 0.014 (0.035 FDR-corrected), Cohen’s d = 0.44; up vs not: t = −0.24, p-value = 0.59 (0.59 FDR-corrected), Cohen’s d =0.045; down vs. not: t = −2.09, p-value = 0.023 (0.035 FDR-corrected), Cohen’s d = 0.39). These behavioral results indicate that TMR differently altered the fate of the motor memory traces depending on the phase of SO during which reactivation was applied. Unexpectedly though, only performance on the down-reactivated sequences differ from the not-reactivated ones.

**Figure 2:**
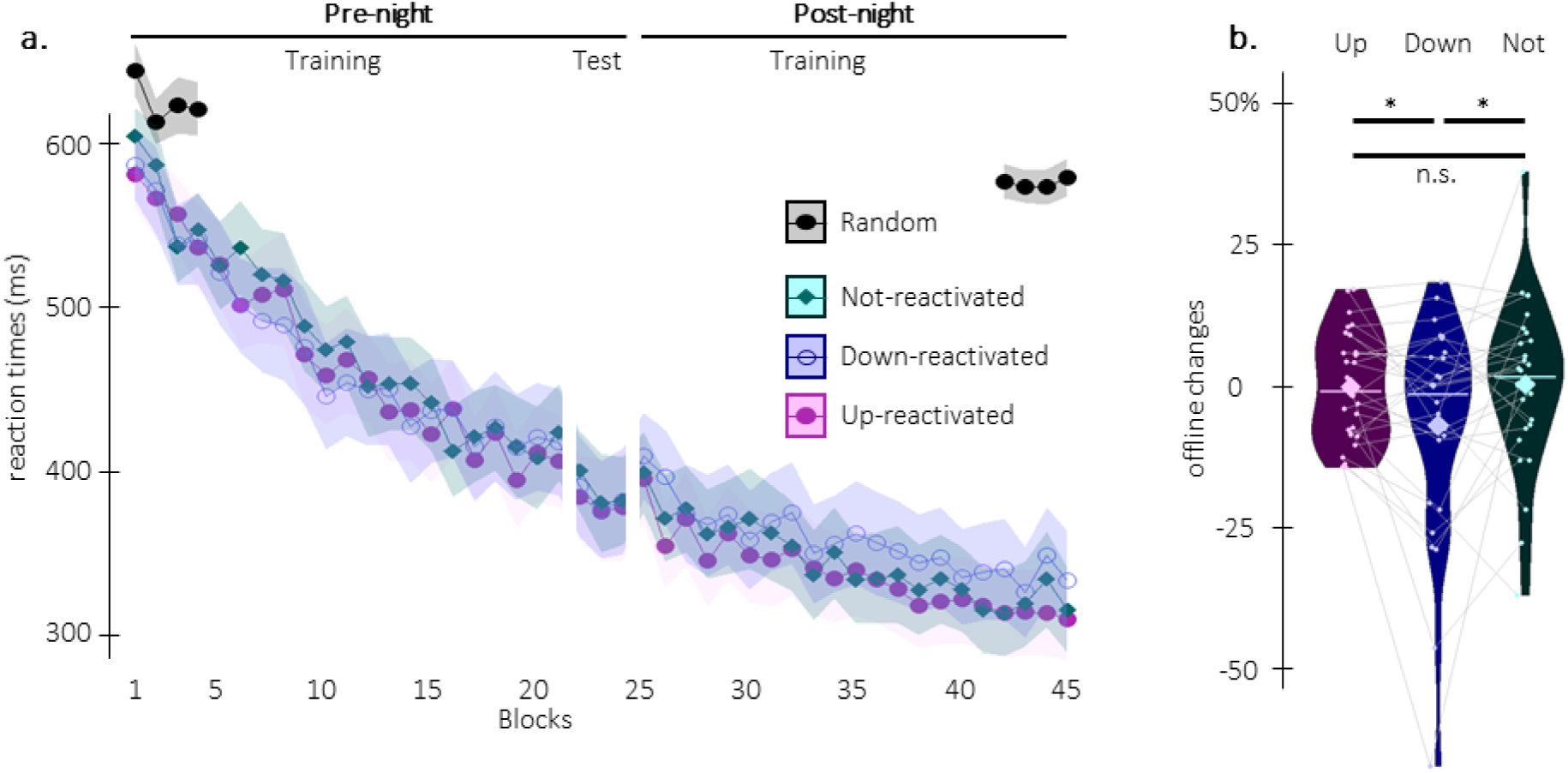
Behavioral results. a. Performance speed. Grand average across participants (n = 28) of median reaction time in ms plotted as a function of blocks of practice during the pre- and post-night sessions (+/- standard error in shaded regions) for the up-reactivated (magenta circles), the down-reactivated (blue empty circles), and the not-reactivated (green diamonds) sequences and for the random serial reaction time task performed at the start and end of the experiment (black overlay, the random task assessed baseline performance and sequence-specific learning, see methods and supplements for corresponding results). Note that a short break is introduced between the training and test runs during the pre-night session in order to minimize the confounding effect of fatigue on end-of-training performance (see methods). The three motor sequences were learned to a similar extent during the pre-night session (see supplemental results). b. Offline changes in performance speed (% change between the average of the three blocks of pre-night test and the first three blocks of post-night training) averaged across participants for the up-reactivated (magenta), the down-reactivated (blue) and the not-reactivated (green) sequences. Results show a main effect of Condition (*: p-value < 0.05) whereby offline changes in performance were greater for the up- and not-reactivated as compared to the down-reactivated sequence (note that improvement in performance from pre- to post-night sessions is reflected by a positive change). Violin plots: median (horizontal bar), mean (diamond), the shape of the violin plots depicts the kernel density estimate of the data. Colored points represent individual data, jittered in arbitrary distances on the x-axis within the respective violin plot to increase perceptibility. For each individual, performance on the different conditions are connected with a line between violin plots. n.s: non-significant.

### 2.2. SO-up-phase TMR enhances both SO amplitude and sigma oscillations

EEG data collected during the TMR episode were analyzed to test whether (the phase of the) stimulation modulated the characteristics of the SOs and spindle/sigma oscillations, two electrophysiological markers critically involved in motor memory consolidation ^24^ and reactivation ^14^ during sleep.

To examine the effect of stimulation on SO characteristics, we computed - for each of the 6 EEG channels (Fz, Cz, Pz, Oz, C3, C4) - event-related potentials locked to the trough of the (i) up-stimulated SOs (detected online on Fpz during up-stimulation intervals), (ii) down-stimulated SOs (detected online on Fpz during down-stimulation intervals) and (iii) not-stimulated SOs (detected offline on Fpz during epochs free of stimulation, see Figure 1b for a depiction of stimulation epochs and Figure S1a in supplemental information for channel level data). In this analysis, cluster-based permutations identified clusters on the basis of temporal and spatial (channel) adjacency (see methods). Results indicated two significant spatio-temporal clusters in which the phase of the stimulation specifically influenced SO amplitude. Specifically, up-stimulated – as compared to down-stimulated - SOs showed (i) greater amplitude around the peak of the SO in a spatial cluster including all electrodes except Oz and (ii) deeper deflection post-peak in a spatial cluster including frontal and central electrodes (up vs. down, Figure 3a). Figure 3a-1 depicts the grand-average of the SOs (superimposed on a time frequency representation of the difference in power modulation, see below) for up and down conditions in which the horizontal black lines represent the significant temporal cluster (see Figure 3a-3 for zoomed inset; Figure 3a-4: positive cluster p-value = 0.0040; Cohen’s d = 0.67 and its topography also showing the spatial dimension of the cluster, i.e., electrodes included in the significant cluster (*); Figure 3a-5: negative cluster p-value = 0.0040; Cohen’s d = −0.60 and its topography also showing electrodes included in the cluster (*)).

**Figure 3:**
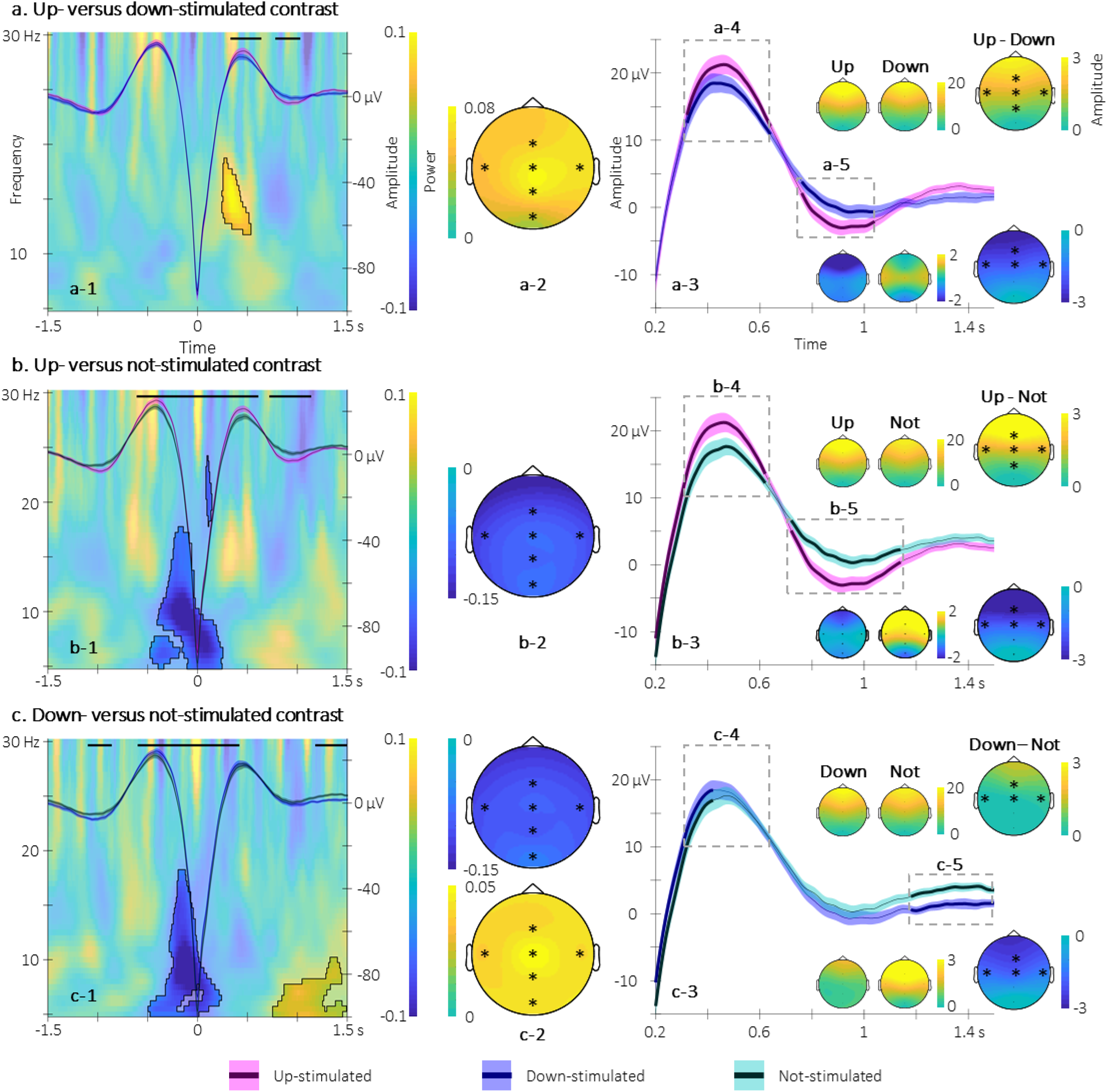
Electrophysiological results. Participants’ sleep was recorded using a 6-channel EEG montage during the night following learning. a. Up- vs down-stimulated contrasts. a-1. Time-frequency representation (TFR) of the difference in power modulation illustrated at Cz around the trough of the up- and down-stimulated SOs on which the grand-average of the SOs illustrated on Fz is super-imposed (magenta(blue): up(down)-stimulated SO). Black lines represent the adjacent time points of the significant spatio-temporal clusters showing a difference in SO amplitude between the two trough-locked ERPs (see a-4 and a-5 for the spatial dimension of the clusters and Figure S1b in supplemental information for channel level cluster depiction). Results show that the up-stimulated SO presented greater amplitude at their peak (from 0.33 to 0.64 sec post-trough) followed by a deeper deflection (from 0.78 to 1.03 sec post-trough). Further, the area highlighted in the TFR represents the adjacent time-frequency points of the significant spatio-temporal-frequency cluster showing a difference in power between conditions. Sigma power nested in the ascending phase of the up-stimulated SOs was greater than for the down-stimulated SOs (from 0.25 to 0.4 sec post-trough and from 12 to 17 Hz; and see a2 for the spatial dimension of the cluster as well as Figure S1b in supplemental information for channel level cluster depiction). a-2. Topography of the significant sigma power modulation. (*) represents the electrodes included in the significant spatio-temporal-frequency cluster. a-3. Zoom on the trough-locked SO peak and deflection at Fz (same color code as a-1) showing the significant differences in amplitude between up and down conditions (see text). a-4 and a-5. Topography of the significant differences in amplitude at the trough-locked SO peak (a-4) and deflection (a-5). (*) represents the electrodes included in the significant spatio-temporal clusters. b. Up- vs not-stimulated contrasts. b-1. Same as a-1 for the up- and the not-stimulated trough-locked SO ERP (magenta and green, respectively) and power modulation. Results show that the amplitude of the up-stimulated SOs was greater than the not-stimulated SOs from −0.61 to 0.61 sec while it reversed from 0.72 to 1.14 sec relative to the trough onset. Power in the 5-17.5 Hz frequency range was lower in the up- as compared to the not-stimulated condition in the descending phase of the SOs (from −0.45 to 0.24 sec relative to the SO trough). b-2. Topography of the significant cluster (between 7-12 Hz and −0.15-0 s time-frequency range) showing that the significant cluster includes all electrodes (*). b-3. Zoom on the trough-locked SO peak and deflection showing the significant differences in amplitude between up and not conditions (see text). b-4 and b-5. Topography of the significant differences in SO amplitude in the peak and deflection time-window defined by the significant clusters highlighted in the up- vs down-stimulated contrast. (*) represents the electrodes included in the significant cluster. c. Down- vs not-stimulated contrasts. c-1. Same as a-1 for the down- and the not-stimulated trough-locked SO ERP (blue and green, respectively) and power modulation. Results show that the amplitude of the down-stimulated SOs was greater than the not-stimulated SOs from −0.60 to 0.42 sec while it reversed from 1.18 to 1.50 sec relative to the trough onset. Power was lower in the 5-18 Hz frequency range in the down- as compared to the not-stimulated condition during the descending phase of SOs (from −0.49 to 0.24 sec relative to the SO trough) and greater in the 5-10 Hz from 0.76 to 1.50 s. c-2. Topography of the negative significant cluster (7-12 Hz and −0.15-0.08 s time-frequency range) and the positive significant cluster (5-8 Hz and 0.8-1.25 s time-frequency range). The significant cluster includes all electrodes (*). c-4 and c-5. Topography of the significant differences in SO amplitude in the peak time-window defined by the significant cluster highlighted in the up- vs down-stimulated contrast and the deflection time-window defined by the significant cluster (1.18-1.50 s post-trough). The significant cluster includes all electrodes except Oz and Pz (*).

**Figure 4:**
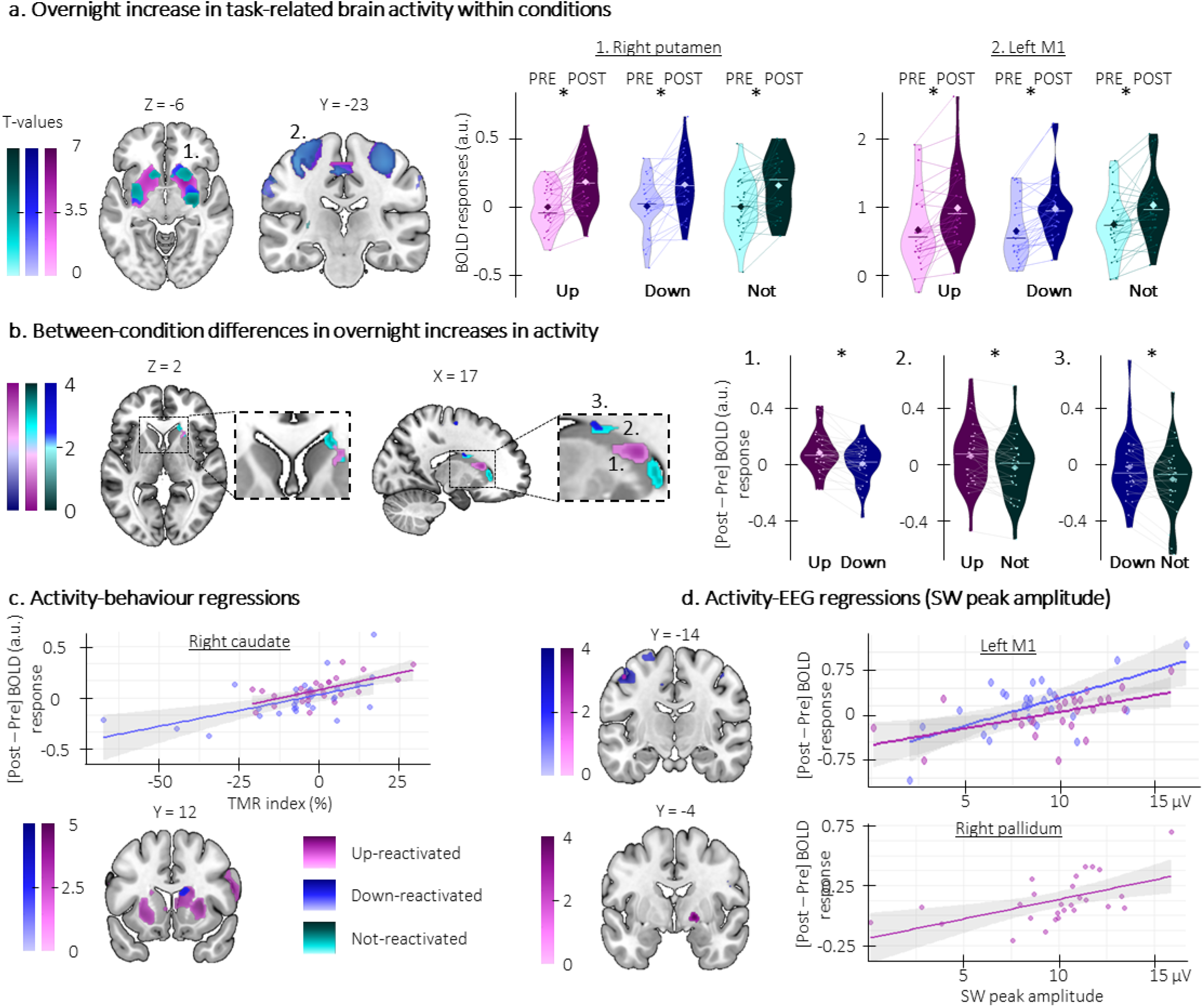
Phase-specific modulation of task-related cortico-striatal activity. a. Overnight increase in activity within condition. Brain activity increased overnight in a set of cortico-striatal regions for up- (Left M1: x = −38, y = −20, z = 52, p_SVC_ <0.001; right putamen: x = 24, y = 18, z = −10, p_SVC_ < 0.001), down- (Left M1: x = −38, y = −24, z = 54, p_SVC_ <0.001; right putamen: x = 26, y = −8, z = −4, p_SVC_ = 0.001), and not-reactivated sequences (Left M1: x = −38, y = −26, z =56, p_SVC_ = 0.004; right putamen: x =28, y = −12, z = −6, p_SVC_ = 0.002). Violin plots represent BOLD responses extracted from clusters overlapping between conditions (1. Right putamen: x = 26, y = −8, z = −4; 2. Left M1: x = −38, y = −20, z = 52). b. Overnight increase in activity between conditions. The overnight increase in striatal (right caudate) activity was greater for the up- as compared to the down-reactivated sequence, which in turn was greater than for the not-reactivated sequence. Violin plots represent the difference in BOLD responses extracted from the activation peaks in the post- versus pre-night sessions (1. up vs. down: x = 20, y = 18, z = 12, p_SVC_ = 0.009; 2. up vs. not: x = 18, y = 28, z =4, p_SVC_ = 0.02; 3. down vs. not: x = 16, y = −2, z = 26, p_SVC_ = 0.016). c. Brain activity-behavior regressions. The overnight increase in striato-motor activity was positively related to the TMR index for both the up- (right caudate: x = 20, y = 18, z = 12, p_SVC_ = 0.005) and down-reactivated sequences (right caudate: x = 16, y = −2, z = 26, p_SVC_ = 0.024) such that the greater the increase in brain activity (i.e., the more positive value on the y-axis), the greater the performance improvement on the reactivated as compared to the not-reactivated sequence (i.e., the more positive TMR index on the x-axis). d. Brain activity-EEG regressions. The overnight increase in activity in the motor cortex (<u>top panel</u>) and the basal ganglia (<u>bottom panel</u>) was related to the SO peak amplitude in the up- (left M1: x = −46, y = −16, z = 48, p_SVC_ = 0.055; right pallidum: x = 20, y = −2, z = −6, p_SVC_ = 0.045) and down-stimulated (left M1: x = −44, y = −16, z = 52, p_SVC_ = 0.012) conditions such that the greater the SO peak amplitude during the night (x-axis), the greater the overnight increase in activity in these regions (y-axis). *: significant after small volume correction (SVC) correction. Activations maps are displayed on a T1-weighted template image with a threshold of p < 0.005 uncorrected. a.u.: arbitrary units. Violin plots: median (horizontal bar), mean (diamond), the shape of the violin plots depicts the kernel density estimate of the data. M1: primary motor cortex.

**Figure 5:**
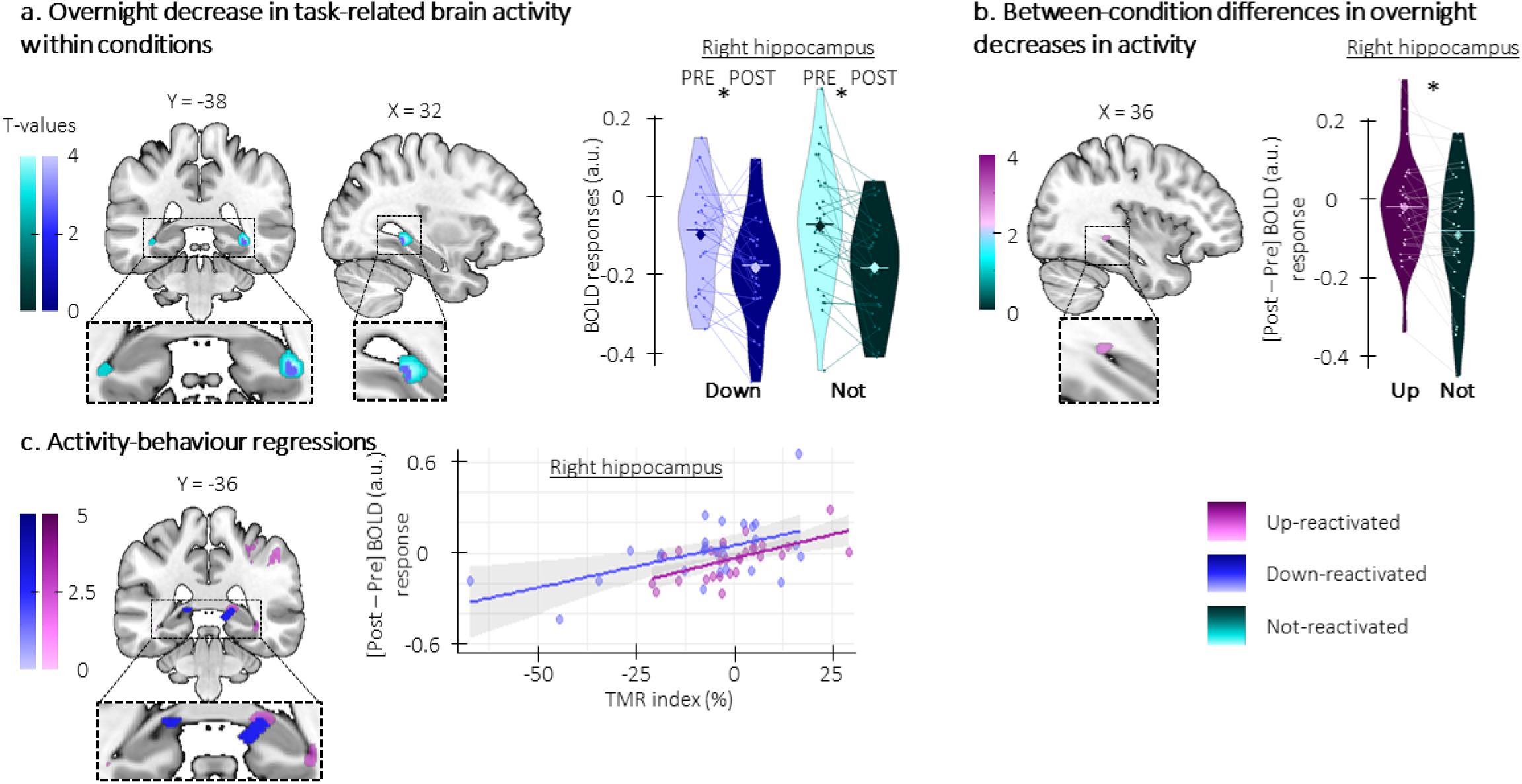
Phase-specific modulation of task-related hippocampal activity. a. Overnight decrease in activity within condition. Brain activity decreased overnight in the hippocampus for both down- (right hippocampus: x = 32, y = −38, z = −6, p_SVC_ = 0.044) and not-stimulated sequences (right hippocampus: x = 34, y = −38, z = −6, p_SVC_ = 0.005). Violin plots represent BOLD responses extracted from clusters overlapping between conditions (x = 32, y = −38, z = −6). b. Overnight decrease in activity between conditions. The overnight decrease in hippocampal activity was greater for the not-reactivated sequence as compared to the up-reactivated sequence (right hippocampus: x = 36, y = −36, z = −4, p_SVC_ = 0.028). Violin plots represent the difference in BOLD responses extracted from the activation peaks in the post- versus pre-night sessions. c. Brain activity-behavior regressions. The overnight increase in hippocampal activity was positively related to the TMR index for both up- (right hippocampus: x = 36, y = −38, z = −8, p_SVC_ = 0.005) and down-reactivated sequences (right hippocampus: x = 20, y = −34, z = 4, p_SVC_ = 0.041) such that the lower the increase in brain activity (y-axis), the lower the performance improvement on the reactivated as compared to the not-reactivated sequence (x-axis). *: significant after SVC correction. Activations maps are displayed on a T1-weighted template image with a threshold of p < 0.005 uncorrected. a.u.: arbitrary units. Violin plots: median (horizontal bar), mean (diamond), the shape of the violin plots depicts the kernel density estimate of the data.

Similar results - albeit on larger time windows - were observed when comparing both up- and down-stimulated versus not-stimulated SOs (up vs. not, Figure 3b-1, horizontal black lines and Figure 3b-3 for zoomed inset; Figure 3b-4: positive cluster p-value = 0.0020, Cohen’s d = 1.12 and its topography; Figure 3b-5; negative cluster p-value = 0.0060, Cohen’s d = −0.70, its topography and electrodes included in cluster; down vs. not, Figure 3c-1, horizontal black lines and Figure 3c-3 for zoomed inset; Figure 3c-4: positive cluster p-value = 0.0020, Cohen’s d = 1.20, its topography and electrodes included in cluster; Figure 3c-5: negative cluster p-value = 0.0040, Cohen’s d = −0.57 and its topography) but that these effects were more pronounced during up- as compared to down-stimulation as shown in Figure 3a.

Note that analogous results were also observed using SO density metrics extracted from the stimulated and not-stimulated blocks (see Figure S2a in supplemental information showing greater density during up-stimulated intervals as compared to down-stimulated and not-stimulated intervals).

To investigate the effect of stimulation on oscillatory brain activity (and sigma oscillations in particular), we performed time-frequency analyses locked to the trough of the stimulated and not-stimulated SOs on each EEG channel. Here, cluster-based permutation analyses identified clusters on the basis of temporal, frequency and spatial adjacency (see methods). Results indicated one significant spatio-temporal-frequency cluster in which sigma power was greater in the ascending phase of the up-stimulated SOs as compared to the down-stimulated SOs on all electrodes (up vs. down: cluster p-value = 0.0080; Cohen’s d = 0.66; Figure 3a-1 for time-frequency representation and a-2 for a display of the topography of this difference and of the electrodes included in the cluster (*)). Note that oscillatory activity in the 5-18 Hz frequency range was lower in the descending phase of both up- and down-stimulated – as compared to not-stimulated – SOs on all electrodes (up vs. not: cluster p-value = 0.002, Cohen’s d = −1.30, Figure 3b-1 and b-2 for topography; down vs. not: cluster p-value = 0.002, Cohen’s d =−0.94, Figure 3c-1 and c-2 for topography). Power in lower frequencies (5-10 Hz) was greater for the down, compared to the not, -stimulated conditions from 0.8 to 1.5 s post SO trough in a cluster including all electrodes (down vs. not: cluster p-value = 0.0020; Cohen’s d = 0.90; Figure 3c-1 and c-2).

Analyses based on sleep spindle events detected from the stimulated and not-stimulated blocks show that spindle frequency and amplitude were unaffected by the stimulation while spindle density was lower during both up- and down-stimulated as compared to not-stimulated blocks, irrespective of the stimulation condition (see Figure S2b-d in supplemental information).

Altogether, these results indicate that up-phase, as compared to down-phase, CL-TMR resulted in enhanced SO density and amplitude as well as a stronger sigma power during the ascending phase of the SO. In contrast, oscillatory activity including the sigma band was decreased during the descending phase of the up- and down-stimulated, as compared to the not-stimulated, SOs and overall spindle density was lower under stimulation, irrespective of its phase.

### 2.3. Phase-specific modulations of task-related hippocampal and striatal activity are related to the effect of TMR on motor performance

Brain imaging data were acquired during task practice before and after the night of stimulation (see Table S1 and Figure S3 for brain activity elicited by task practice during initial learning). We first examined whether task-related brain activity *increased* from the pre-night to the post-night practice sessions within each condition. Results showed, for all conditions, a strong overnight increase in task-related brain activity in a set of striato-cortical regions including the putamen and the primary motor cortex (Figure 4a and see Table S2-1 for a complete list of activations). Interestingly, the overnight increase in striatal activity was greater for the up-reactivated sequence as compared to the down-reactivated sequence, which in turn was greater than for the not-reactivated sequence (Figure 4b; see Table S2-2 of the supplemental information for details).

Importantly, the between-session increase in striato-motor activity reported above was correlated with the TMR index (i.e., the difference in offline changes in performance between the reactivated vs. the not-reactivated sequences) for both the up- and down-reactivated sequences (Figure 4c; see Table S2-3). We also performed exploratory analyses to probe the link between the sleep EEG features showing the phase-specific modulation described above (i.e., SO amplitude and sigma power at the peak of the SO) and the between-session changes in brain activity. These analyses did not reveal any correlation between brain activity and sigma power but they showed that the overnight increase in activity in the basal-ganglia and the motor cortex was related to greater SO peak amplitude in the up and down conditions (Figure 4d; see Table S2-4 of the supplemental information). Altogether, the regression analyses indicate that greater overnight increase in striato-motor activity is related to both greater SO amplitude during the post-learning night and greater overnight gains in motor performance in both up and down conditions. Interestingly, despite condition differences in overnight changes in brain activity (Figure 4b), SO amplitude (Figure 3a) and motor performance (Figure 2), the phase of the stimulation did not alter the relationship between these differences.

Next, we examined whether task-related brain activity *decreased* between the pre- and post-night practice sessions. Results showed that hippocampal activity decreased overnight for both the down- and the not-reactivated sequences while no significant changes were observed in the up condition (Figure 5a; see Table S2-1 of the supplemental information). The decrease in hippocampal activity observed for the not-reactivated sequence was greater than for the up-reactivated sequence (Figure 5b; see Table S2-2 of the supplemental information). Importantly, the between-session changes in hippocampal activity reported above were correlated with the TMR index for both the up- and down-reactivated sequences such that greater overnight decrease in activity was related to poorer performance (Figure 5c; see Table S2-3 of the supplemental information). Finally, we did not observe any relationships between EEG features and changes in hippocampal activity. Overall, these results suggest that up-stimulation prevented the overnight decrease in hippocampal activity observed in the other conditions, the amplitude of which is related to poorer performance.

Altogether, these results show that the amplitude of the changes in brain activity occurring in striato-hippocampo-motor areas as a result of the consolidation process were modulated by the phase of the SO during which TMR was applied. Importantly, the magnitude of these changes was related to SO characteristics and changes in motor performance, both metrics that also showed a phase-specific modulation of amplitude.

### 2.4. Phase-specific modulations of connectivity in striato-hippocampo-motor networks are related to the effect of TMR on motor performance

We examined whether stimulation modulated task-related connectivity patterns in the brain regions showing phase-specific modulation of activity described above (i.e., the hippocampus and the striatum, see methods and Table S3).

We observed an overnight decrease in hippocampo-motor connectivity for the up-reactivated sequence which was greater than for the not-reactivated sequence (Figure 6a and 6b, see Tables S3-1.3.1 and - 2.3.2 of the supplemental information). Moreover, an overnight decrease in striato-cortical (Figure S4a, right panel and Tables S3-3.2.1 of the supplemental information) and striato-hippocampal connectivity was related to a greater TMR index for the up-reactivated sequence (Figure 6c, see Tables S3-3.3.1 of the supplemental information). This suggests that the beneficial effect of up-stimulation on performance was paralleled by more segregation of task-relevant brain regions within their functional network, i.e., by a decrease in connectivity between these brain areas (which was also paralleled by an overall increase in activity within these brain regions, see above).

**Figure 6:**
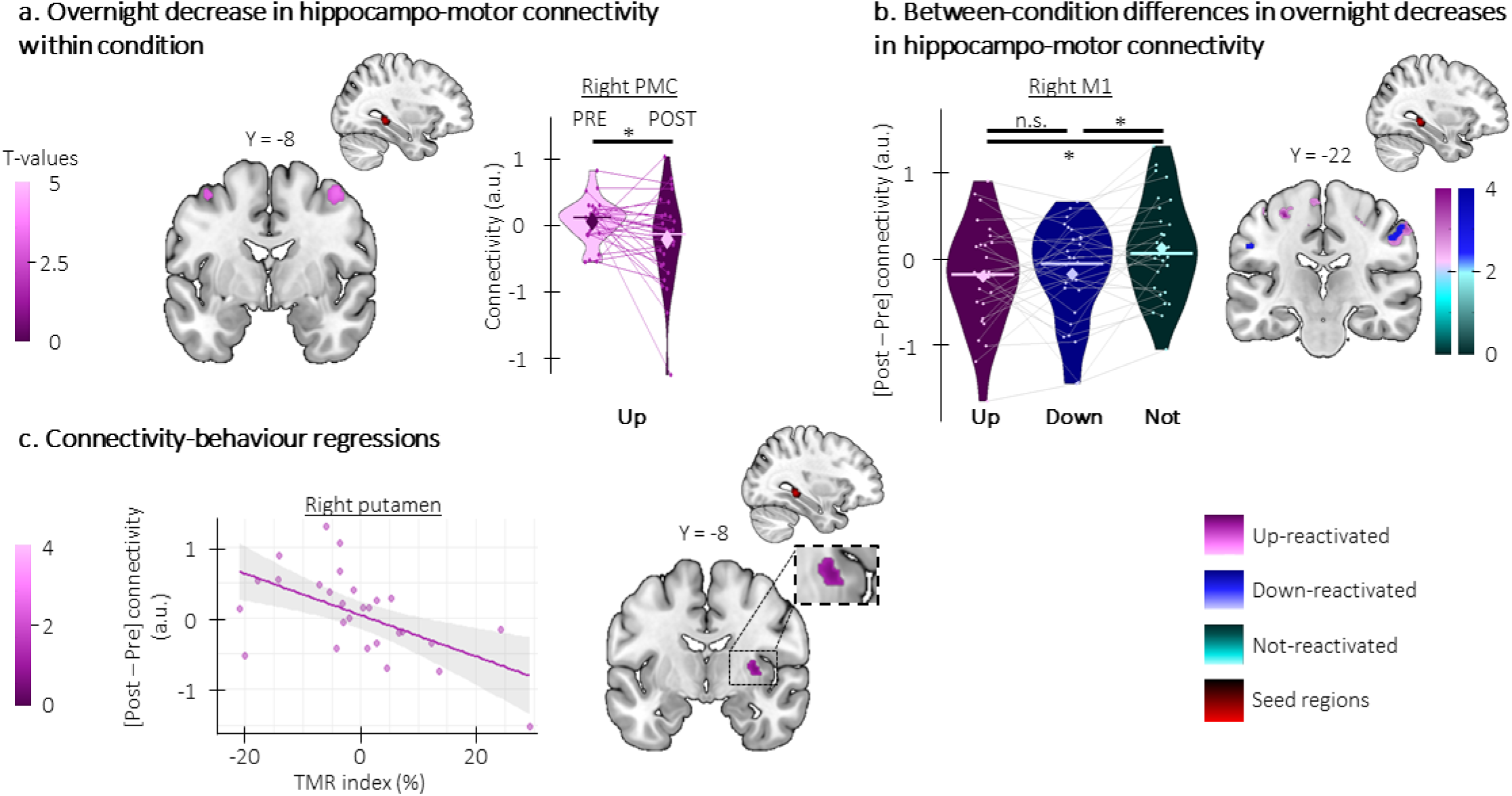
Phase-specific modulation of task-related striato-hippocampo-cortical connectivity. a. Overnight decrease in hippocampo-motor connectivity within condition. Hippocampo-motor connectivity decreased from pre- to post-night sessions for the up-reactivated condition (hippocampus-right PMC: x = 46, y = −8, z = 54, p_SVC_ = 0.035). Violin plots represent BOLD responses extracted from this cluster. b. Overnight decrease in hippocampo-motor connectivity between conditions. The hippocampo-motor connectivity overnight decrease was greater in the up- (hippocampus-right M1: x = 58, y = −22, z = 46, p_SVC_ = 0.033) and the down-reactivated sequences (hippocampus-right M1: x = 54, y = −22, z = 46, p_SVC_ = 0.046) as compared to the not-reactivated sequence. Violin plots represent BOLD responses around a common significant voxel (x = 52, y = −24, z = 40). c. Brain connectivity-behavior regressions. The overnight decrease in striato-hippocampal (hippocampus-right putamen: x = 32, y = −8, z = 4, p_SVC_ = 0.027) connectivity was negatively correlated with the TMR index such that the greater the decrease in connectivity (y-axis), the greater the performance improvement on the up-reactivated as compared to the not-reactivated sequence (x-axis). M1: primary motor cortex. *: significant after SVC correction). Activations maps are displayed on a T1-weighted template image with a threshold of p < 0.005 uncorrected. a.u.: arbitrary units. Violin plots: median (horizontal bar), mean (diamond), the shape of the violin plots depicts the kernel density estimate of the data.

In the down-reactivated condition, there was an overnight increase in striato-motor connectivity (Figure 7a; see Tables S3-1.2.2 of the supplemental information) that was greater than for the up-reactivated sequence (Figure 7b; see Tables S3-2.2.1 of the supplemental information). Interestingly, we observed an overall negative relationship between overnight increases in connectivity in hippocampo-striato-motor networks and sleep features such that lower SO amplitude and sigma power were related to greater overnight increases in connectivity (striato-hippocampal connectivity-sigma power: see Figure 7c, left panel and Tables S3-4.1.2; striato-motor connectivity-SO amplitude: see Figure 7c, right panel; and Tables S3-4.4.2 and 4.5.2; hippo-motor connectivity-SO amplitude: see Figure S4b and Tables S3-4.6.2; and striato-hippocampal connectivity-SO amplitude: see Figure S4c; and Tables S3-4.6.2). These results suggest that the reduced amplitude of sleep features observed under down- (as compared to up-) stimulation was presumably related to compensatory overnight increases in connectivity in hippocampo-striato-motor networks. Importantly, these overnight increases in connectivity were differently related to behavior depending on the networks examined. Specifically, the overnight increase in striato-motor connectivity was related to poorer TMR index (Figure 7d, left panel; see Tables S3-3.2.2 of the supplemental information) while the increase in striato-hippocampal connectivity was related to greater TMR index (Figure 7d, right panel, see Tables S3-3.1.2 of the supplemental information). These findings suggest that increases in connectivity in striato-motor networks were ineffective to compensate for the negative effect of down-stimulation on performance while increases in connectivity between the striatum and the hippocampus were related to greater performance improvement.

**Figure 7:**
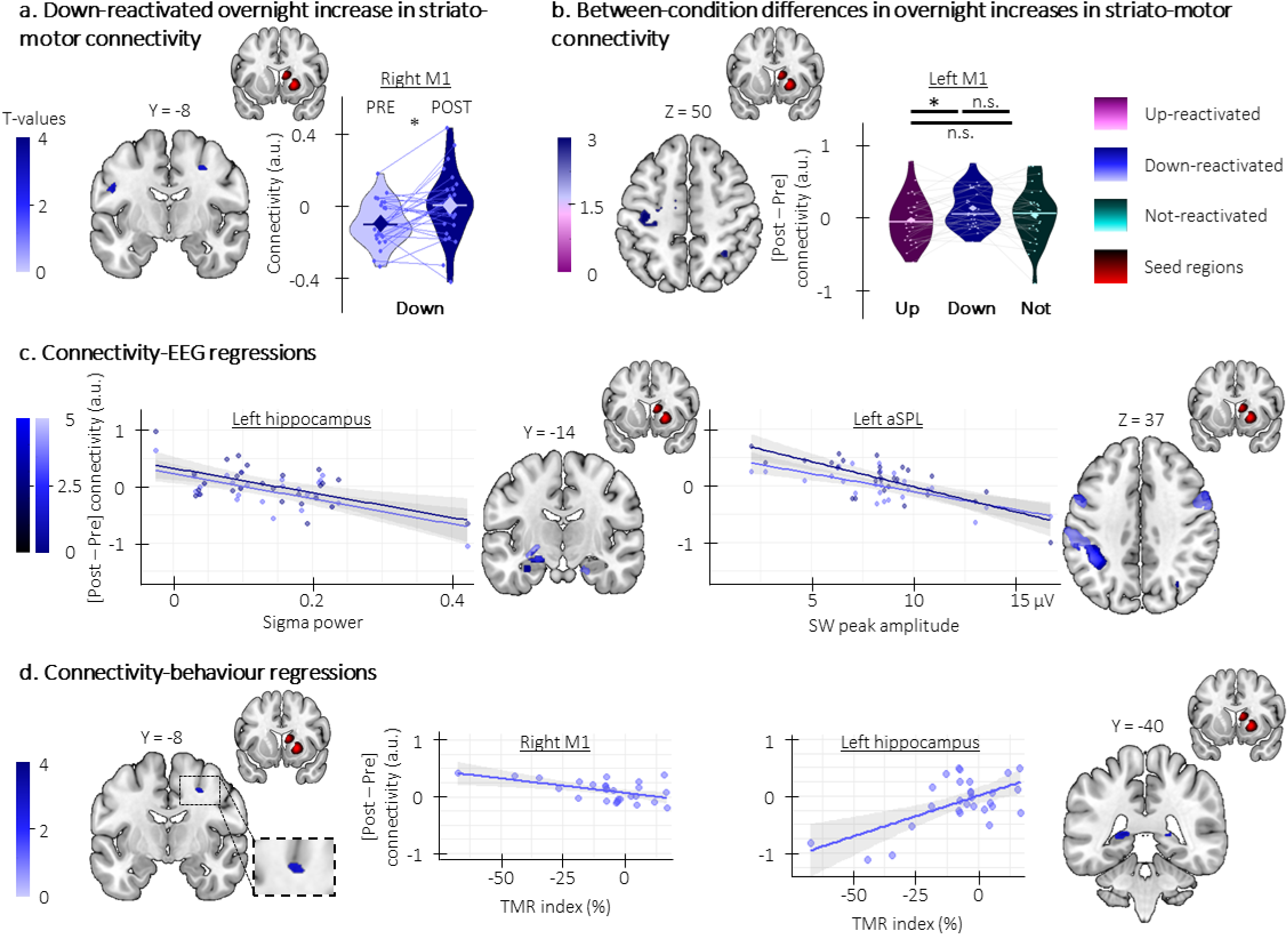
Down-stimulation modulation of striato-hippocampo-cortical connectivity. a. Overnight increase in striato-motor connectivity. Striato-motor connectivity increased from pre- to post-night session for the down-reactivated condition (Putamen-right M1: x = 28, y = −8, z = 44, p_SVC_ = 0.038). Violin plots represent BOLD responses extracted from the cluster. b. Overnight increase in striato-motor connectivity between conditions. The overnight increase in striato-motor connectivity was greater in the down- as compared to the up-reactivated sequence (Putamen-left M1: x = −32, y = −20, z = 50, p_SVC_ = 0.02). Violin plots represent BOLD responses extracted from the cluster. c. Brain connectivity-EEG regressions. The overnight decrease in striato-hippocampal (<u>left</u> <u>panel</u>, Caudate-left hippocampus (pale blue): x = −24, y = −14, z = −8, p_SVC_ = 0.005; Putamen-left hippocampus (dark blue): x = −18, y = −10, z = −8, p_SVC_ = 0.017) and striato-motor (<u>right panel</u>, Caudate-left aSPL (pale blue): x = −44, y = −44, z = 38, p_SVC_ = 0.003; Putamen-left aSPL (dark blue): x = −38, y = −42, z = 36, p_SVC_ < 0.001) connectivity were correlated with the sigma power and SO peak amplitude respectively such that the lower the SO peak amplitude and sigma power during the night (y-axis), the greater the overnight increase in connectivity (x-axis). d. Brain connectivity-behavior regressions. The overnight increase in striato-motor connectivity was related to the TMR index (<u>left panel</u>, Putamen-right M1: x = 26, y = −8, z = 44, p_SVC_ = 0.025) such that the greater the increase in brain connectivity (y-axis), the lower the performance improvement on the down-reactivated as compared to the not-reactivated sequence (x-axis). In contrast, the overnight increase in striato-hippocampal connectivity was positively related to the TMR index (<u>right panel,</u> caudate-left hippocampus: x = −16, y = −40, z = 6, p_SVC_ = 0.014) such that the greater the increase in brain connectivity (y-axis), the greater the performance improvement on the down-reactivated as compared to the not-reactivated sequence (x-axis). M1: primary motor cortex; *: significant after SVC correction. Activations maps are displayed on a T1-weighted template image with a threshold of p < 0.005 uncorrected. a.u.: arbitrary units. Violin plots: median (horizontal bar), mean (diamond), the shape of the violin plots depicts the kernel density estimate of the data.

In sum, the connectivity results indicate an overall decrease in hippocampal and striatal connectivity after up-stimulation that was related to better performance. In contrast, down-stimulation resulted in an overall increase in connectivity in hippocampo-striato-motor networks that was related to the lower amplitude of sleep features during down-stimulation. Interestingly, these overnight increases in connectivity were differently related to performance improvement suggesting that different networks may play distinct roles to compensate for the reduced plasticity induced by down-stimulation during sleep.

## 3. Discussion

The goal of this pre-registered study was to combine targeted memory reactivation (TMR) and closed-loop (CL) stimulation approaches to (i) test whether reactivating motor memories at the up (as compared to the down)-phase of slow oscillations (SOs) during post-learning sleep could enhance motor memory consolidation; and to (ii) provide a comprehensive characterization of the underlying neurophysiological processes using sleep EEG and task-related fMRI. As hypothesized, overnight changes in performance were greater for motor sequences reactivated at the up-, as compared to the down-phase of the SO. Unexpectedly though, only performance on the down-reactivated sequence differ from the not-reactivated one. Electrophysiological data showed that up-stimulated SOs were of higher amplitude and presented greater peak-nested sigma power (spindle frequency band) than down-stimulated SOs. Brain imaging data collected during task practice indicated that the practice of up-, as compared to down-reactivated sequences, resulted in greater activity in striato-motor areas, greater maintenance in hippocampal activity, and decreased connectivity in these networks. Importantly, these modulations in brain responses were related to the up-TMR-induced increase in SO amplitude and improvement in performance. In contrast, down-stimulation resulted in a lower increase in striato-motor activity that was paralleled by significant increases in connectivity in striato-hippocampo-motor networks, and both were related to the lower amplitude of sleep features during down-stimulation. Interestingly, the overnight increases in connectivity observed after down-stimulation were related to better (striato-hippocampus) or worse (striato-motor) performance on the down-reactivated sequence.

Our behavioral results indicate that TMR applied at the up-phase of the SO resulted in greater gains in motor performance than when administered at the down-phase of the SO. These phase-specific effects are in line with previous studies in which acoustic stimulations delivered in a closed-loop fashion at the up-phase (or during the down-to-up transition) of the SO have been shown to enhance declarative memory consolidation ^17,20,21,25–27^ (but see ^26,28^ for null effects). We are only aware of one study using closed-loop acoustic stimulation in the motor memory domain and results showed no benefit of SO up-stimulation on motor performance ^29^. The discrepancy between this recent research and our findings is unclear but we speculate that methodological differences between studies, such as time afforded in NREM sleep (nap vs. night paradigm) or stimulation phase (380ms post-trough vs. peak), might have contributed to these inconsistencies. Another notable difference is that the sounds used in the current research were memory cues. In line with the current findings, the few studies using SO-closed-loop stimulation with memory cues (i.e., CL-TMR) show an up-phase stimulation benefit on declarative memory consolidation as compared to no-reactivation ^18^ or down-stimulation ^19^. Overall, our results concord with this earlier research and suggest that reactivating motor memories at the up-, as compared to the down-phase, of the slow oscillations during post-learning sleep benefits motor memory consolidation. Interestingly, our results also indicate that down-reactivated sequences presented significantly worse performance as compared to both up- and not-reactivated sequences. One could therefore argue that down-stimulation actively disrupted the motor memory consolidation process. These findings contradict earlier studies showing no specific effect of down-, as compared to no-stimulation, on memory ^17–19^. Based on this earlier research and evidence that down-phases of SOs are silent phases of neuronal inactivity ^30^, we speculate that our results are driven by the nature of our paradigm rather than by an active disruptive effect of down-stimulation on memory consolidation. Specifically, it is possible that, due to our within-subject design, overall acoustic stimulation during post-learning sleep might also have boosted performance on the control (not-reactivated) condition. This could also explain the lack of difference between up and not conditions in the current study. While this is hypothetical, it is in line with previous work using similar within-subject design showing no difference in performance between the up- and not-reactivated conditions ^19^.

Our electrophysiological data show phase-specific modulations of sleep oscillations such that up-stimulated SOs exhibited higher amplitude and presented greater sigma power nested at their peak as compared to down-stimulated SOs. Together with sleep spindles and sharp wave ripples, slow oscillations are part of the three cardinal NREM sleep oscillations playing a critical role in memory consolidation during sleep ^1,6,31^. SO up-states have received particular attention as they are known to host heightened excitability ^30^ It is thought that prolonging SO up-phase with stimulation increases the probability of neuronal ensembles to fire together and strengthen the memory traces encoded in these networks ^32^. Accordingly, prior research has used experimental interventions to target plasticity processes during this window and, in turn, influence memory consolidation during sleep. In line with our findings, this earlier research has collectively shown that stimulating SO at the up-phase of the slow-oscillation enhanced the amplitude of ongoing SOs ^17–21,25–27,29,33^, sigma oscillation power during the ascending phase of the SO ^18–20^ and memory consolidation ^17–21,26,27^. Interestingly, the time-locking of the sigma burst to the up-phase of the SO has been shown to predict a positive outcome of consolidation ^14,34^. Altogether, our findings generally concord with earlier experimental work and with models suggesting that the synchronous neural firing orchestrated by the SOs leads to a higher probability of downstream synchronous neural firing favoring the occurrence of higher frequency oscillation bursts such as sleep spindles that are beneficial for memory consolidation ^35^.

Our brain imaging data indicate that up-stimulation resulted in an overnight increase in task-related striato-motor activity and a maintenance of hippocampal activity as compared to the down- and no-stimulation conditions. The involvement of striato- and hippocampo-cortical networks in motor sequence learning and memory consolidation is well documented. Specifically, task-related striato-motor activity generally increases with consolidation and during later stages of learning ^e.g.,22,36^. Hippocampal activity during both learning and delayed retests has also been associated to successful (sleep-related) motor memory consolidation ^24,37–40^. The present data therefore suggest that up-stimulation further strengthened the modulation of brain activity that is usually observed as a result of the spontaneous motor memory consolidation process. Interestingly, our results also show that the modulations of hippocampal and striato-motor activity reported above were related to both the overnight performance improvement and the enhancement of sleep features. These findings generally concord with a series of earlier correlational studies. First, they are in line with an acoustic stimulation study showing that up-phase stimulation-induced increases in SO amplitude were correlated with greater hippocampal activity during subsequent (post-sleep) declarative learning ^33^. They also concur with earlier studies in the motor memory domain showing a positive relationship between spindle-locked striatal and hippocampal reactivations during post-learning sleep and offline gains in motor performance ^41^. They are also consistent with the findings of a previous TMR study showing that the increased striatal and hippocampal activity observed for reactivated (as compared to not-reactivated) motor sequences was related to the time spent in NREM sleep (during which TMR was applied) ^12^. Based on the evidence reviewed above and the current data, we suggest that the up-stimulation-induced increase in SO amplitude and sigma power during post-learning sleep facilitated hippocampal and striatal reactivations which in turn resulted in greater activity in these networks during retest and better motor performance. It is worth noting that similar modulations of brain activity - albeit of lower amplitude - were observed for down-reactivated sequences. Interestingly, the relationship between the modulation in brain responses in the down condition and both performance and sleep features was similar as for the up condition but scaled down in terms of amplitude (i.e., lower changes in brain activity, lower performance gains and lower amplitude of sleep features). These results suggest that plasticity processes induced by down-stimulation during sleep were presumably reduced as compared to up-stimulation and therefore less favorable for reactivation processes to take place during post-learning sleep. This is in line with previous research suggesting that the efficacy of TMR follows a (down-to-up) gradient rather than an “all-or-none principle” ^19^. Here, we speculate that such reduced plasticity resulted in lower activity in hippocampal and striatal areas during retest and poorer motor performance. Altogether, these brain imaging results further corroborate the hypothesis that down-stimulation did not actively disrupt motor memory consolidation but rather failed to potentiate ongoing consolidation processes.

Our brain connectivity analyses revealed an overnight decrease in hippocampo-striato-cortical connectivity in the up-condition which was related to better performance. These observations stand in contrast with earlier reports of increased connectivity within striato-motor networks ^42^ and between the striatum and the hippocampus ^24,43^ as a result of spontaneous motor memory consolidation. They are also inconsistent with earlier TMR studies showing greater connectivity in these networks during the practice of reactivated as compared to not-reactivated motor sequences ^12,44^. We speculate that the decrease in connectivity observed after up stimulation reflects an advanced stage of consolidation in which brain regions are more segregated within their functional networks due to a decreased need for long-range integration as a result of consolidation ^1,32,45,46^. In contrast to these observations, connectivity analyses in the down condition revealed an overnight increase in connectivity in these networks which was negatively correlated to sleep features such that lower SO amplitude and sigma oscillation power were related to greater connectivity increase. Together with the brain activation results showing lower modulation of brain activity after down-stimulation, it is tempting to speculate that these large increases in connectivity reflect compensatory processes. Importantly, the increases in connectivity observed for the down-reactivated sequences were differently related to changes in performance depending on the network involved. Specifically, the overnight increase in striato-hippocampal connectivity was associated to better performance while increases in striato-motor connectivity were related to worse performance. We speculate that connectivity within these different networks reflects different processes. On the one hand, connectivity between the striatum and the hippocampus during initial motor learning has been proposed to reflect the integration of different tasks components (motor vs. spatial) that is necessary to optimize motor behavior ^22,24^. On the other hand, increases in striato-motor connectivity occur at later stages of learning for the automatization of the motoric component of the task ^36,47^. We argue that the sub-optimal plasticity process under down-stimulation slowed down consolidation such that greater interaction between the striatal and hippocampal systems - usually observed during initial learning - was beneficial for performance at retest while greater connectivity in non-optimally consolidated motor networks failed to optimize behavior. This remains, however, speculative.

In summary, this study showed that motor memories reactivated during the up-phase of slow oscillations exhibited superior consolidation compared to memories reactivated at the down-phase. Our results suggest that the phase-specific beneficial effect of slow-oscillation closed-loop TMR on consolidation was related to phase-specific modulations of activity and connectivity in task-relevant networks including the striatum, the hippocampus and the motor cortex which were also associated to phase-specific alterations of the characteristics of sleep EEG features involved in plasticity processes. Altogether, this study not only highlights the promise of up-phase CL-TMR to impact motor memory consolidation but also sheds light on the complex interplay between sleep oscillations, task-related brain activity and connectivity patterns, and motor performance.

## 4. Material and Methods

This study was pre-registered in the Open Science Framework (https://osf.io/). Our pre-registration document outlined our hypotheses and intended analysis plan as well as the statistical models used to test our a priori hypotheses (available at https://osf.io/dpu6z). Whenever an analysis presented in the current paper was not pre-registered, it is referred to as exploratory. Additionally, to increase transparency, any deviation from the pre-registration is marked in the methods section with a (#) symbol and listed in Table S4 of the supplemental information together with a justification for the change.

### 4.1. Participants

Young healthy volunteers (23.7 yo ranging from 18 to 30) were recruited to participate in the proposed research project and received a monetary compensation for their time and effort. Every participant gave written informed consent before participating in this research protocol, which was approved by the local Ethics Committee (B322201525025) and was conducted according to the declaration of Helsinki (2013). Inclusion criteria were: 1) no previous extensive training of dexterous finger movements via playing a musical instrument or as a professional typist, 2) free of medical, neurological, psychological, or psychiatric conditions, including depression ^48^ and anxiety ^49^, 3) no indications of self-reported abnormal sleep ^50^, 4) free of psychoactive and sleep-influencing medications, 5) eligible for MR measurements, and 6) right-handed ^51^. None of the participants were shift-workers or did a trans-meridian trip in the month preceding the study.

We performed a power analysis based on our previous study investigating auditory TMR in an open-loop paradigm ^14^. This power analysis was performed with the G*Power software ^52^. The partial η² was calculated based on our behavioral main effect showing a significant effect of condition (reactivated vs. control) on offline changes in performance speed and was transformed to an effect size f (η² = 0.15; f = 0.42). The correlation coefficient calculated between the offline changes in performance speed in the reactivated and the control conditions was 0.66, but due to the different nature of the design (e.g., 3 sequences instead of 2, 5-element sequences instead of 8), we set the average correlation coefficient between repeated measures at r = 0.5 for a more conservative power calculation. Finally, the sphericity correction was set to 0.5 since the primary factors of interest in our design had 3 levels (up-reactivated, down-reactivated and not-reactivated). The primary contrast of interest is a reactivation condition main effect on offline changes in performance speed tested with a one-way rmANOVA. The required sample size for a 95% power is 27 at an alpha error probability of 0.05. In total, 34 (age range 18 – 30 yo) participants completed the study. Two participants were excluded for experimental error (i.e., one participant because they were erroneously enrolled despite being left handed and the other one because of technical errors at the scanner) and one for excessive movement in the scanner (see MRI section below). The remaining 31 participants were included in closed-loop stimulation analyses (i.e., detection accuracy) but only 27 participants presented a complete dataset (behavior, sleep EEG and MRI data). In three participants, only the sleep EEG data was analyzed as behavioral and MRI data of the post-night session was corrupted due to experimental error (sleep EEG analyses, N = 30). For one participant, only MRI and behavioral data were analyzed due to an EEG recording default (Behavioral and MRI analyses, N = 28). Also note that for five participants, the pre-night Psychomotor Vigilance Task data was overwritten due to experimental error. Participant’s characteristics are reported in Table S5 in the supplemental information.

### 4.2. General design

In this full within-subject design presented in Figure 1, participants were first invited, in the evening, for a habituation night during which they completed a full night of sleep monitored with polysomnography (PSG). Roughly a week later, participants returned to complete the experimental session. Between these two visits, each participant followed a constant sleep/wake schedule (according to their own rhythm +/- 1h) during at least 3 days before the experimental session (compliance assessed with sleep diaries and wrist actigraphy, ActiGraph wGT3X-BT, Pensacola, FL). Sleep quality and quantity for the night preceding the experiment was assessed with the St. Mary’s sleep questionnaire ^53^ (sleep data are presented in Table S5).

During the experiment, volunteers participated in two fMRI sessions referred to as pre-night (around 9 pm) and post-night (around 8.30 am) sessions which took place at the MRI facility as well as an experimental night (EEG) session which took place in the sleep lab between the two fMRI sessions (between 11pm-7am). At the beginning of each fMRI session, vigilance was measured objectively and subjectively using the Psychomotor Vigilance Task (PVT) ^54^ and Stanford Sleepiness Scale (SSS) ^55^, respectively (vigilance data are reported in Table S5). The pre-night MRI session started with a resting state scan (RS1). Then participants performed two motor tasks: (i) the random version of the serial reaction time task (SRTT, see below for details) to measure baseline performance / general motor execution (not scanned) and (ii) the training and a short post-training test on the sequential SRTT probing motor sequence learning (both scanned). During the sequential SRTT, participants learned to perform three different motor sequences. Each sequence was associated to a different sound. Two of the three sounds associated to the learned material were replayed - using auditory closed-loop stimulation - on the peak (up-state) or the trough (down-state) of slow oscillations detected online during the subsequent sleep episode (see Polysomnography and TMR section for detection algorithm details). The other sequence served as a no-reactivation control condition. The combination between the 3 reactivation conditions (referred to as up-reactivated, down-reactivated and not-reactivated conditions), the 3 sounds and the 3 sequences was randomized across participants. After task completion, a post-task RS scan (RS2) and a structural scan were acquired. Following this first MRI session, participants were transferred to the sleep lab. During the experimental night, participants’ brain activity was recorded with PSG and CL-TMR was applied during the NREM2-3 stages of the first 3 hours of sleep (i.e. 3 hours from the first stimulation). The post-night session took place at the MRI facility and started with a RS (RS3) that was followed by the retest on the sequential SRTT. The session was concluded with the last RS (RS4) and the random SRTT (not scanned). Note that the RS data are not reported in the present manuscript.

### 4.3. Stimuli and tasks

#### 4.3.1. Motor Task

A bimanual serial reaction time task (SRTT) ^56,57^ was implemented in Matlab Psychophysics Toolbox version 3 ^58^. During the task, eight squares were presented on the screen, each corresponding to one of the eight keys on a specialized MR-compatible keyboard and to one of the 8 fingers of both hands excluding the thumbs. The color of the outline of the squares alternated between red and green, indicating rest and practice blocks, respectively. During the practice block, participants had to press as quickly as possible the key corresponding to the location of a green filled square that appeared on the screen. After a response, the next square changed to green following either a pseudo-random (see below for details) or sequential order depending on the task variant. After 20 presses, the practice block automatically turned into a rest block and the outline of the squares changed from green to red. The rest interval was 10s.

In the sequential version of the SRTT, participants learned 3 different 5-element sequences (sequence A: 4 7 2 8 3, sequence B: 1 6 3 5 2, and sequence C: 7 3 8 4 6, where 1 is the left little finger and 8 is the right little finger) that were pseudo-randomly assigned to the three conditions. Sequence practice was organized by block (one sequence practiced per block) and each sequence was repeated 4 times per block (20 key presses per block). The order of the sequences was fixed within participants but pseudo-randomized across participants. The participants were explicitly instructed that the visual cues would appear following a sequential pattern and that there would be three different motor sequences to perform. The pre-night training session consisted of 63 practice blocks (21 blocks per sequence) immediately followed by a post-training test of 9 practice blocks (3 blocks per sequence). This test was performed in order to minimize the confounding effect of fatigue on end-of-training performance ^59^. The post-night session consisted of 63 practice blocks (21 blocks per sequence). During all task practice sessions, three different 100-ms auditory cues (see below) pseudo-randomly assigned to sequences A, B, or C were played before the beginning of each sequence.

For the pseudo-random version of the SRTT, four blocks composed of 12 sequences were created by randomly selecting 5 keys out of 8 at each iteration (60 key presses per block). For both version of the SRTT, participants were instructed to keep their fingers still and look at the squares on the screen during the rest blocks and to respond as quickly and as accurately as possible to the visual cues during the practice blocks.

#### 4.3.2. Acoustic stimulation

The same three different 100-ms sounds used our previous research ^14,60^ were pseudo-randomly assigned to the three conditions (up-reactivated, down-reactivated, and not-reactivated), for each participant. The three synthesized sounds consisted of (1) a tonal harmonic complex created by summing a sinusoidal wave with a fundamental frequency of 543 Hz and 11 harmonics with linearly decreasing amplitude (i.e. the amplitude of successive harmonics is multiplied by values spaced evenly between 1 and 0.1); (2) a white noise band-passed between 100-1000 Hz; and (3) a tonal harmonic complex created with a fundamental frequency of 1480 Hz and 11 harmonics with linearly increasing amplitude (i.e. the amplitude of successive harmonics is multiplied by values spaced evenly between 0.1 and 1). A 10-ms linear ramp was applied to the onset and offset of the sound files so as to avoid earphone clicks. Before the start of the training session, a dummy MRI acquisition was launched to adjust the volume of the three different sounds. Sounds were played via MR-compatible, electrostatic headphones (MR-Confon, Magdeburg, Germany). An experimenter adjusted the volume of the sounds until the participant reported they could hear it above and beyond the scanner noise but still comfortably. The sound level determined for each of the three sounds was then used during task practice. During the reactivation session taking place during the experimental night in the sleep lab, sounds were played via ER3C air tube insert earphones (Etymotic Research). Before turning the light offs for the night, auditory detection thresholds were determined by performing a transformed 1-down 1-up procedure ^61,62^ separately for each of the three sounds. Subsequently, the sound pressure level was set to 2db above the individual auditory threshold, thus limiting the risk of awakening during the night. The three sounds were then presented to the participants at the intensity mentioned above to confirm that they could hear them distinctively. Before the start of the night episode, participants were instructed that they may or may not receive auditory stimulations during the night.

### 4.4. EEG data acquisition and closed-loop TMR

Both habituation and experimental nights were monitored with a digital sleep recorder (V-Amp, Brain Products, Gilching, Germany; bandwidth: DC to Nyquist frequency) and were digitized at a sampling rate of 1000 Hz. Standard electroencephalographic (EEG) recordings were made from Fz, C3, Cz, C4, Pz, Oz, A1 and A2 according to the international 10-20 system (note that Fz, Pz and Oz were omitted during habituation). A2 was used as the recording reference and A1 as a supplemental individual EEG channel. An electrode placed on the middle of the forehead was used as the recording ground. Bipolar vertical and horizontal eye movements (electrooculogram: EOG) were recorded from electrodes placed above and below the right eye and on the outer canthus of both eyes, respectively. Bipolar submental electromyogram (EMG) recordings were made from the chin. Electrical noise was filtered using a 50 Hz notch.

The CL-TMR device required another set of electrodes for which the signal was recorded from FPz (ground and reference electrodes placed behind the right ear). During the experimental night, an experienced researcher performed online visual scoring of the polysomnography (PSG) data in order to detect NREM2-3 sleep. When these stages were reached, the phase detection algorithm was launched (see below). The auditory stimulation was presented in a blocked design with 3-min long intervals that alternated between up- and down-SO detection/stimulation. Each stimulation block was separated by a 1-minute silent period (Figure 1B). The stimulation was manually stopped when the experimenter detected REM sleep, NREM1 or wakefulness. The CL-TMR ended 3 hours after the first stimulation was sent (about 2 sleep cycles). The sounds associated to the up(down)-reactivated sequence was then played on the peak(trough) of the SOs within these alternating blocks. The algorithm for online SO detection consisted of a two-step process for trough and peak detection. For down-detection, a fast-moving average filter was employed with a window of 50 samples and a trough was detected when signal went below a specific threshold adapted for biological sex according to ^23^ and of −41µV in females and 39.5µV in males. For up-detection, the peak of a SO was identified when, in addition to the criterion described for trough detection above, peak-to-peak signal amplitude reached 77 µV in females and 74 µV in males. Importantly, as the trough detection relied on less criteria than peak detection, the likelihood to detect trough was higher than the one for peak. To address this issue, a secondary filter was implemented to look backwards and validate the initial detections. This algorithm assessed whether the detected events corresponded to true slow oscillations. By tracking the true positive count for both conditions, the algorithm dynamically adjusted its detection strategy. If the count indicated an imbalance, with more down- than up-detections, the algorithm temporarily paused during down-detection intervals, allowing up-detection to catch up. The ultimate goal was to achieve balanced stimulation between conditions (Table S5 in supplemental information).

### 4.5. fMRI data acquisition

MRI data were acquired on a Philips Achieva 3.0T MRI system equipped with a 32-channel head coil. Task-related fMRI data were acquired during the training and overnight retest sessions using an ascending gradient EPI pulse sequence for T2*-weighted images (TR = 2000 ms; TE = 29.8 ms; multiband factor 2; flip angle = 90°; 54 transverse slices; slice thickness = 2.5 mm; interslice gap = 0.2 mm; voxel size = 2.5 × 2.5 × 2.5 mm^3^; field of view = 210 × 210 × 145.6 mm^3^; matrix = 84 × 82) for each participant (max. 1200 dynamic scans). Resting-state fMRI data were also collected prior and immediately after the training and overnight retest sessions with the same EPI sequence as above (data not reported here). Additionally, field maps (TR = 1500 ms; TE = 3.5 ms; flip angle = 90°; 42 transverse slices; slice thickness = 3.75 mm; interslice gap = 0 mm; voxel size = 3.75 × 3.75 × 3.75 mm^3^; field of view = 240 × 240 × 157.5 mm^3^; matrix = 64 × 64) were collected immediately before the SRTT Training and Retest together with three sets of EPI images using reversed phase-encoding polarity (TR = 2000 ms; TE = 29.8 ms; multiband factor 2; flip angle = 90°; 54 transverse slices; slice thickness = 2.5 mm; interslice gap = 0.2 mm; voxel size = 2.5 × 2.5 × 2.5 mm^3^; field of view = 210 × 210 × 145.6 mm^3^; matrix = 84 × 82, 6 dynamic scans). Note that these sequences were not included in the final analysis pipeline^#^ (see #1 in Table S4 of the supplemental information). High-resolution T1-weighted structural images were acquired with a MPRAGE sequence (TR = 9.5 ms, TE = 4.6 ms, TI = 858.1 ms, FA = 9°, 160 slices, FoV = 250 × 250 mm2, matrix size = 256 × 256 × 160, voxel size = 0.98 × 0.98 × 1.20 mm3) for each participant.

### 4.6. Analyses

#### 4.6.1. Behavioral data

##### 4.6.1.1. Preprocessing

*Motor performance* on both the random and sequential SRTT was measured in terms of speed (median of correct response time RT, in ms) and accuracy (% of correct responses, with a trial classified as “correct” if the key pressed by the participants matches the visual cue) for each block of practice. Note that correct trials were excluded from the analyses if they were outlier trials based on John Tukey’s method of leveraging the Interquartile Range^#^ (5.1% of the correct trials were outliers, see #2 in Table S4 of the supplemental information). Consistent with our pre-registration, our primary analyses focused on performance speed (but see Figure S5 in the supplemental information for results related to the accuracy).

The *offline changes in performance* on the sequential SRTT were computed as the relative change in speed between the end of the training of the pre-night session (namely during the 3 blocks of the pre-night test) and the beginning of the post-night session (3 first blocks of practice) separately for the up-reactivated, the down-reactivated, and the not-reactivated sequences. A positive offline change in performance therefore reflects an increase of absolute performance from the pre-night test to the post-night test. Additionally, we computed a TMR index which consisted of the difference in offline gains in performance between up-reactivated and not-reactivated sequences (TMR index_up_) and down-reactivated and not-reactivated sequences (TMR index_down_), separately. A positive TMR index reflects higher offline changes in performance for the reactivated sequences as compared to the not-reactivated, control, one.

##### 4.6.1.2. Statistics

Behavioral statistical analyses were performed with the open-source software R ^64,65^. Statistical tests were considered significant for p < 0.05. When necessary, corrections for multiple comparisons were conducted with the False Discovery Rate ^66^ (FDR) procedure within each family of hypothesis tests. Greenhouse-Geisser corrections was applied in the event of the violation of sphericity. F and t statistics and corrected p-values were therefore reported for ANOVAs and student tests, respectively. Effect sizes are reported using *Cohen’s d* for Student t-tests and *η²* for rmANOVAs using G*power ^52^.

We describe in the supplemental information the negative control analyses that were collectively designed to verify that the pattern of behavioral results emerged from our experimental manipulation on motor memory consolidation processes rather than from the various potential confounding factors listed below. First, we tested whether vigilance during each behavioral session was similar using a one-way rmANOVAs on both the median RTs of the PVT and the Stanford Sleepiness Scale scores with Session as two-level factor (pre- and post-night, see Table S5). Second, we tested whether the three movement sequences were learned to the same extent during the pre-night session using two-way rmANOVAs on performance speed and accuracy measures with sequence (A vs. B vs. C) and blocks (21 for training and 3 for post-training test) as within-subject factors (Figure S6 in the supplemental information). Third, we performed the same analysis using condition (up-reactivated vs. down-reactivated vs. not-reactivated) – as opposed to sequence (A, B and C) - and blocks (21 for training and 3 for post-training test) as within-subject factors (Figure 2a). Last, to highlight that improvement in movement speed was specific to the learned sequences as opposed to general improvement of motor execution, we computed the *overall performance change* for both the sequential SRTT (first 4 blocks of the pre-night training vs. 4 last blocks of post-night training collapsed across sequences) and the pseudo-random version of the SRTT (4 blocks pre-night session vs. 4 blocks post-night session).

In our confirmatory analysis, we tested whether offline changes in performance on the sequential SRTT differed between reactivation conditions after a night of sleep. To do so, a one-way rmANOVA was performed on the offline changes in performance speed (main text) and accuracy (supplemental information and Figure S5) with Condition (up- vs. down- vs. not-reactivated) as within-subject factor. Post-hoc analysis on the 3 possible pair comparisons were performed using Student t tests and FDR correction was applied ^66^.

#### 4.6.2. Electrophysiological data

##### 4.6.2.1. Offline sleep scoring

Offline sleep scoring was performed by a certified sleep technologist - blind to the stimulation periods - according to criteria defined in the guidelines from the American Academy of Sleep Medicine ^67,68^ using the software SleepWorks (version 9.1.0 Build 3042, Natus Medical Incorporated, Ontario, Canada). Data were visually scored in 30 s epochs and band pass filters were applied between 0.3 and 35 Hz for EEG signals, 0.3 and 30 Hz for EOG, and 10 and 100 Hz for EMG. A 50 Hz notch filter was also used. Sleep characteristics resulting from the offline sleep scoring as well as the distribution of auditory cues across sleep stages and SO phases are shown in Table S5 of the supplemental information. Briefly, results indicate that participants slept 7.5 hours on average (sleep efficiency: 83.3 %) and that cues were accurately presented in NREM sleep (stimulation accuracy mean: 98.5% (95CI: 97.6 - 99.3); up-reactivated cues: 98.8 % (95CI: 98.1 - 99.4); down-reactivated cues: 98.2% (95CI: 97.2 - 99.3) and at the correct phase (true positive mean: 82.2 % (95CI: 79.6 – 84.8); up-reactivated: 89.5 % (95CI: 87.8 - 91.2); down-reactivated: 74.9 % (95CI: 71.7 - 78.1); see Figure S7 of the supplemental information).

##### 4.6.2.2. Preprocessing

EEG data preprocessing was carried out using functions supplied by the fieldtrip toolbox ^69^. EEG was re-referenced to an average of A1 and A2 and filtered between 0.1-30 Hz. Specifically, data were cleaned by manually screening each 30-sec epoch. Data segments contaminated with muscular activity or eye movements were excluded. Independent component analysis was used to remove cardiac artifacts. We then offline detected the SOs on the FPz channel use for online detection with criteria published in previous research and similar to our online detection ^23,70^. The trough time-sample of each offline-detected and stimulated SO (referred to as true positive in Table S5) was extracted from the reactivation period (i.e. the first three hours of the night) for both the up- and down-stimulation blocks. We also extracted the trough time-sample of each offline-detected SO occurring during the silent intervals (referred to as not-stimulated SO).

##### 4.6.2.3. Event-related analyses

Event-related data analyses included trough-locked potentials and oscillatory activity and were performed with down sampled data (100Hz). Trough-locked responses were obtained by segmenting the data into epochs time-locked to the trough of the SOs offline-detected on FPz (from −2 to 2 sec) for the up-, the down- and the not-stimulated SO and averaged across all trials ^#^ (see #3 in Table S4 of the supplemental information) in each condition separately. The average number of artifact-free trials by condition was of 622.9 [95% CI: 495.1 – 750.6] for the up-, 599.7 [95% CI: 482.4 – 717.1] for the down-, and 664.3 [95% CI: 528.6 – 800.0] for the not-stimulated conditions. To analyze oscillatory activity, we computed Time-Frequency Representations (TFRs) of the power spectra per experimental condition and per channel. To this end, we used an adaptive sliding time window of five cycles length per frequency (Δt = 5/f; 20-ms step size), and estimated power using the Hanning taper/FFT approach between 5 and 30 Hz. Individual TFRs were converted into change of power relative to the entire period around the SO trough (from −2s to 2s relative to trough) ^#^ (see #3 in Table S4 of the supplemental information). Note that statistical analyses were performed on a more conservative 1.5s to 1.5s relative to SO-trough to avoid border effects. Nonparametric CBP tests ^71^ implemented in fieldtrip toolbox were used for both ERP and TF analyses. For both analyses, we used paired t-test between conditions and cluster-based correction (Maris and Oostenveld, 2007) to account for multiple comparisons across time and space for the ERP analyses, and time, frequency and space for the TF analyses. All time-space (ERP analyses) and time-frequency-space (TF analyses) samples whose t values exceeded a threshold of alpha cluster of 0.01 were considered as candidate members of clusters, i.e. samples clustered in connected sets on the basis of time and space adjacencies for ERP analyses and on the basis of time, frequency and space adjacencies for the TF analyses. The sum of t-values within every cluster, that is, the ‘cluster size’, was calculated as test statistics. These cluster sizes were then tested against the distribution of cluster sizes obtained for 500 partitions with randomly assigned conditions within each individual. The clusters were considered significant at *p* < 0.05. For CBP contrast analyses, *Cohen’s d* is reported. Corrections for three comparisons, i.e., p < 0.0083, was conducted with Bonferonni procedure within each family of hypothesis tests^#^ (see #4 in Table S4 of the supplemental information).

Note that event-related phase amplitude coupling analyses were also pre-registered but eventually not performed as redundant with SO-trough locked analyses^#^ (see #5 in Table S4 of the supplemental information).

##### 4.6.2.4. Sleep events detection

Induced sleep spindles and SOs were detected on all EEG channels automatically a posteriori in NREM sleep epochs during the reactivation period by using the YASA open-source Python toolbox ^72^. This analysis on induced events included all detected sleep events in blocks of stimulated and not-stimulated intervals. Preprocessed cleaned data were down-sampled to 500 Hz and were transferred to the python environment. Concerning the spindle detection, the algorithm is inspired from the A7 algorithm described in Lacourse et al. ^73^. The relative power in the spindle frequency band (12–16 Hz) with respect to the total power in the broad-band frequency (1–30 Hz) is estimated based on Short-Time Fourier Transforms with 2-s windows and a 200-ms overlap. Next, the algorithm uses a 300ms window with a step size of 100 ms to compute the moving root mean squared (RMS) of the filtered EEG data in the sigma band. A moving correlation between the broadband signal (1–30 Hz) and the EEG signal filtered in the spindle band is then computed. Sleep spindles are detected when the three following thresholds are reached simultaneously: (i) the relative power in the sigma band (with respect to total power) is above 0.2 (ii) the moving RMS crosses the RMS_mean_ + 1.5 RM_SSD_ threshold and (iii) the moving correlation described is above 0.65. Additionally, detected spindles shorter than 0.5 s or longer than 2 s were discarded. Spindles occurring in different channels within 500ms of each other were assumed to reflect the same spindle. In these cases, the spindles are merged together. Concerning the SO detection, the algorithm used is a custom adaptation from ^70,74^. Specifically, data were filtered between 0.3 and 2 Hz with a FIR filter using a 0.2 Hz transition resulting in a –6 dB points at 0.2 and 2.1 Hz. Then all the negative peaks with an amplitude between –40 and –200 μV and the positive peaks with an amplitude comprised between 10–150 μV are detected in the filtered signal. After sorting identified negative peaks with subsequent positive peaks, a set of logical thresholds are applied to identify the true slow oscillations: (1) duration of the negative peak between 0.3 and 1.5 sec; (2) duration of the positive peak between 0.1 and 1 sec; (3) amplitude of the negative peak between 40 and 300 µV; (4) amplitude of the positive peak between 10 and 200 µV and (5) PTP amplitude between 75 and 500 µV.

We extracted the frequency and the amplitude of spindles as well as the density of both spindles and SO. On these variables of interest, we performed one-way rmANOVAs with condition (events occurring during up- vs. down- vs. not-stimulated intervals) as within-subject factor using the software R ^64,65^ and Greenhouse-Geisser corrections was applied in the event of the violation of sphericity. Statistical tests were considered significant for p < 0.05. When a condition effect was detected, post-hoc analysis on the 3 possible pair comparisons were performed using Student t-test and FDR correction was applied^66^.

#### 4.6.3. fMRI data

Statistical parametric mapping (SPM12; Welcome Department of Imaging Neuroscience, London, UK) was used for the preprocessing of the functional images and the statistical analyses of BOLD data.

##### 4.6.3.1. Preprocessing

Preprocessing included the realignment of the functional time series using rigid body transformations, iteratively optimized to minimize the residual sum of squares between each functional image and the first image of each session separately in a first step and with the across-session mean functional image in a second step (mean of maximum movement in the three dimensions: 1.49 mm (95CI: 0.91 – 2.07) for the pre-night training session and 0.96 mm (95CI: 0.73 – 1.18) for the post-night training session). Movement was considered as excessive when exceeding more than 2 voxels mm in either or the three dimensions for both the pre- and post-night sessions (one individual was excluded from data analyses because of such excessive movement). The pre-processed functional images were then co-registered to the structural T1-image using rigid body transformation optimized to maximize the normalized mutual information between the two images. The anatomical image was segmented into gray matter, white matter, cerebrospinal fluid (CSF), bone, soft tissue and background and the individuals’ forward deformation fields were used for the normalization step. All functional and anatomical images were normalized to the MNI template (resampling size of 2 x 2 x 2 mm). Functional images were spatially smoothed using an isotropic 8 mm fullwidth at half-maximum [FWHM] Gaussian kernel.

##### 4.6.3.2. Activation-based analyses

The analysis of the task-based fMRI data, based on a summary statistics approach, was conducted in two serial steps accounting for intra-individual (fixed effects) and inter-individual (random effects) variance, respectively. Changes in brain regional responses was estimated for each participant with a model including responses to the three motor sequences (up- vs. down- vs. not-reactivated) and their linear modulation by performance speed (median RT on correct key presses per block) for each task run (pre-night training, pre-night test, post-night training). The rest blocks occurring between each block of motor practice served as the baseline condition modeled implicitly in the block design. These regressors consisted of boxcars convolved with the canonical hemodynamic response function. Movement parameters derived from realignment as well as erroneous key presses were included as covariates of no interest. High-pass filtering was implemented in the design matrix using a cutoff period of 128 s to remove slow drifts from the time series. Serial correlations in the fMRI signal was estimated using an autoregressive (order 1) plus white noise model and a restricted maximum likelihood (ReML) algorithm. Linear contrasts were generated at the individual level to test for (1) the main effect of practice (across sequences) and its linear modulation by performance, (2) the main effect of practice for each sequence (up-, down-, and not-reactivated) and (3) the difference in brain responses between sequences (reactivated-up vs. reactivated-down vs. not-reactivated). These contrasts were written within each of the two training runs (pre-night and post-night training) ^#^ (see #6 in Table S4 of the supplemental information for justification) as well as between these runs. The resulting contrast images were further spatially smoothed (Gaussian kernel 6 mm Full Width at Half Maximum (FWHM)). The resulting contrast images were entered in a second level analysis for statistical inference at the group level (one sample t-tests), corresponding to a random effects model accounting for inter-subject variance.

##### 4.6.3.3. Connectivity-based analyses

Task-related functional connectivity was examined using psychophysiological interaction (PPI) analyses. Specifically, we assessed connectivity of three seed regions (right caudate (x = 10, y = 14, z = 12), right putamen (x = 18, y = 12, z = −2), and right hippocampus (x = 32, y = −38, z = −6)) revealed by the univariate analyses and showing a main effect of session across multiple conditions (see Figure S8 in supplemental information). In order to limit the number of PPI analyses, we opted to use right (instead of both right and left) seeds as they showed preferential phase-dependent modulation of activity (see results presented in Table S2-2). For each individual, the first eigenvariate of the signal was extracted using Singular Value Decomposition of the time series across the voxels included in a 10 mm-radius sphere centered on these coordinates. Linear models were generated, at the individual level, with a first regressor representing the practice of the motor sequence (pre- and post-night sessions in each of the three reactivation conditions), a second regressor corresponding to the BOLD signal in the seed and a third regressor representing the interaction between the first (psychological) and second (physiological) regressors. To build this regressor, the underlying neuronal activity was first estimated by a parametric empirical Bayes formulation, combined with the psychological factor, and subsequently convolved with the hemodynamic response function ^75^. The individual linear contrasts testing for the interaction between the psychological and physiological regressors within and between the different runs mentioned above were then further spatially smoothed (Gaussian kernel 6 mm FWHM). The resulting contrast images were entered in a second level analysis for statistical inference at the group level (one sample t-tests), corresponding to a random effects model accounting for inter-subject variance.

##### 4.6.3.4. Regression analyses

We performed regression analysis between the individuals’ brain maps showing between session changes in activity/connectivity within each condition and the individuals’ TMR index (for each condition separately, i.e. TMRindex_up_ and TMRindex_down_). These regressions were performed in a second level analysis for statistical interference at the group level (one sample t-test), corresponding to a random effects model accounting for inter-subject variance. Finally, we performed exploratory regression analyses between the individuals’ brain maps showing between session changes in activity/connectivity within each condition and the EEG sigma power as well as the peak amplitude of the SOs. For these analyses, the significant clusters from the event-related potentials and the oscillatory activity analyses of the up- vs down-stimulated contrasts were used (see Figure 2 in the main manuscript and Figure S1b in the supplemental information). The amplitude and power for each individual were averaged across all channels between 0.32-0.64 sec post-trough (peak amplitude) and between 0.25-0.4 sec post-trough and 12-17 Hz (sigma power).

##### 4.6.3.5. Statistics

The set of voxel values resulting from each second level analysis described above (activation, functional connectivity and regression analyses) constituted maps of the t statistic [SPM(T)], thresholded at p < 0.005 (uncorrected for multiple comparisons). The goal of the fMRI analyses was to examine brain patterns elicited in specific regions of interest (ROIs). The following (bilateral) ROIs were selected a priori based on previous literature describing their critical involvement in motor sequence learning processes ^22,76,77^: the primary motor cortex (M1), the supplementary motor cortex (SMA), the premotor cortex (PMC), the anterior part of the superior parietal lobule (aSPL), the hippocampus, the putamen and the caudate nucleus. These ROIs were defined with the brainnetome atlas as follows. M1 contained the upper limb and hand function regions of Brodmann area (BA) 4. The premotor cortex (PMC) was defined as the dorsal (A6cdl; dorsal PMC) and ventral (A6cvl; ventral PMC) part of BA 6. aSPL was defined to include the rostrocaudal areas of inferior parietal lobel (39rd and 40rd), as well as the intraparietal area 7 (A7ip) and the lateral area 5 of the superior parietal lobe (A5l). The SMA was defined as part A6m of the superior frontal gyris and area 4 of the paracentral lobule (a4ll). The probability maps of these cortical areas were thesholded at 50% for binarization. The hippocampus mask included both rostral and caudal parts of the hippocampus. The caudate mask included both dorsal and ventral parts of the caudate. The putamen mask included both ventromedial and dorsololateral part of the putamen. The probability maps of these subcortical areas were thesholded at 5% for binarization.

Statistical inferences were performed at a threshold of p < 0.05 after family-wise error (FWE) correction for multiple comparisons over small volume (SVC, 10 mm radius) located in the structures of interest reported by published work (see Table S6 in supplemental information). All results reported and discussed in the main text survived SVC.

## Footnotes

1 Note that whenever an analysis presented in the current paper was not pre-registered, it is referred to as exploratory. Additionally, any deviation from the pre-registration is marked in the methods section with a (#) symbol and listed in Table S4 of the supplemental information together with a justification for the change.

