## Supplemental information for "Neurophysiological Correlates of Phase-Specific Enhancement of Motor Memory Consolidation via Slow-Wave Closed-Loop Targeted Memory Reactivation"

CHANNEL LEVEL SLEEP EEG DATA

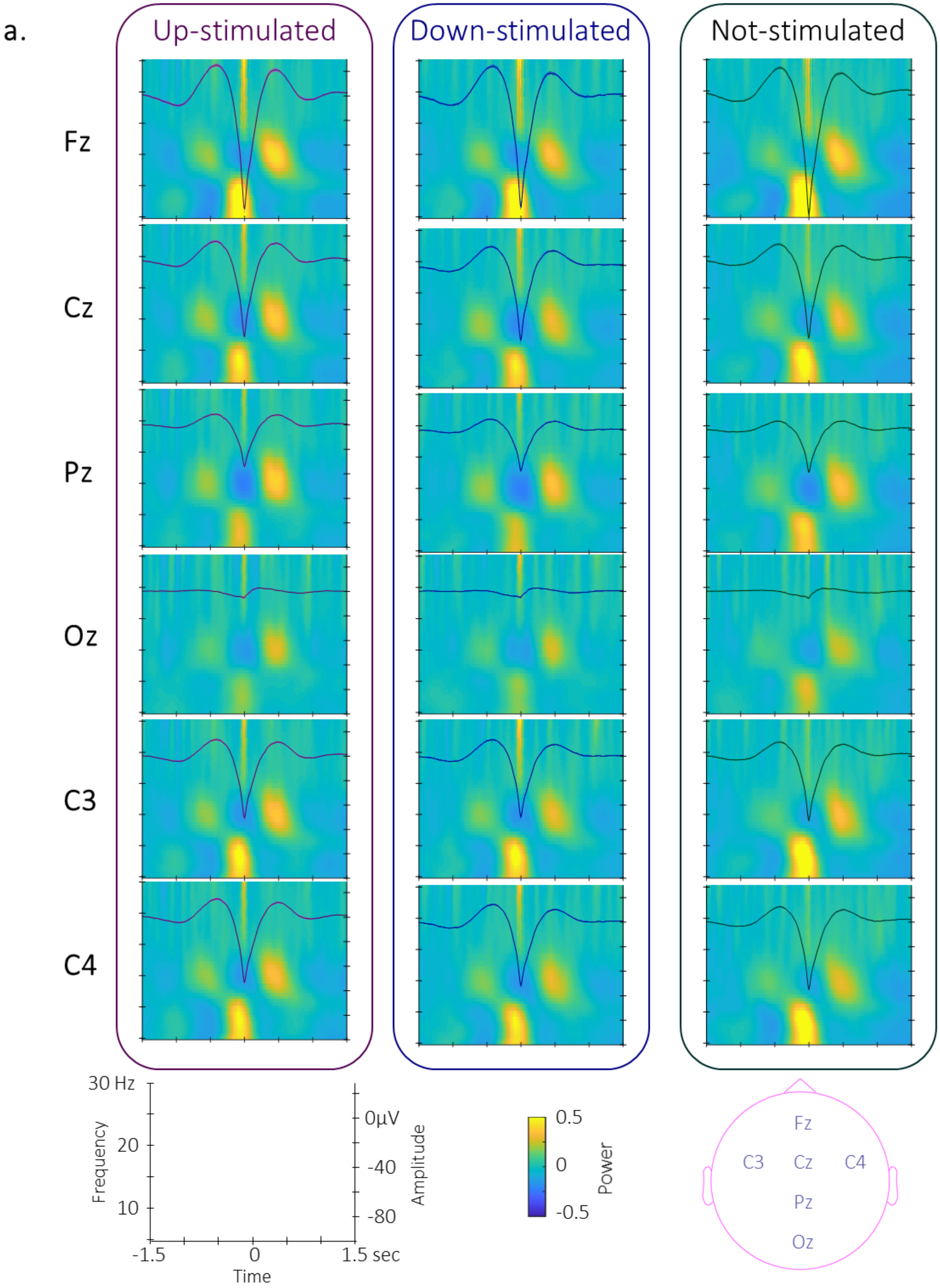

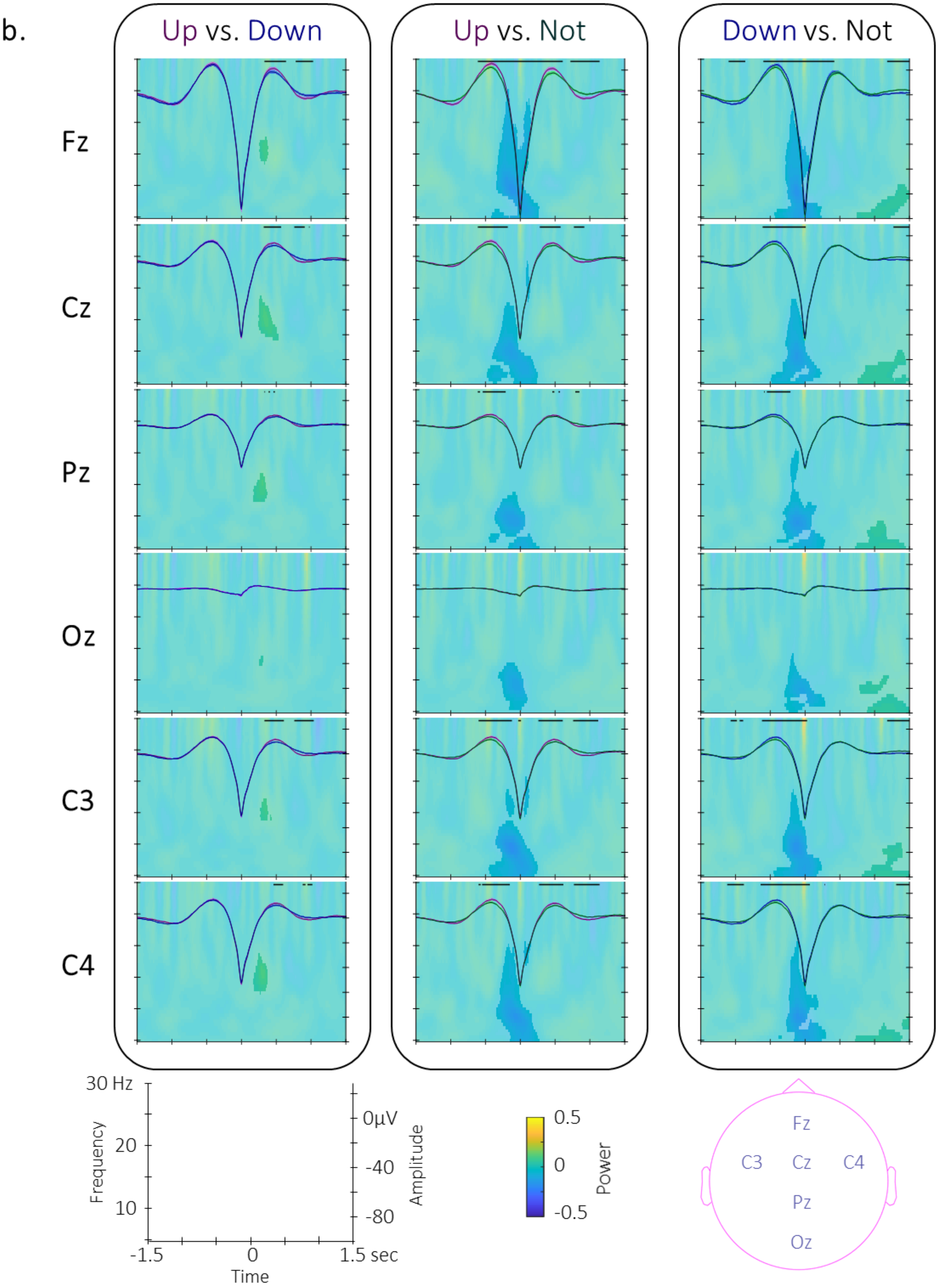
b.
Up vs. Down
Up vs. Not
Down vs. Not
Fz
Cz
Pz
Oz
C3
C4
30 Hz
20
10
Frequency
-1.5
0
1.5 sec
Time
0μV
-40
-80
Amplitude
0.5
0
-0.5
Power
Fz
C3
Cz
C4
Pz
Oz

*Figure S1 (previous page): Channel-level EEG data. a.* *Time-frequency representation (TFR) of the group average time-frequency power per condition on which is super-imposed the group average (+/- standard error in shaded regions) of trough-locked SW (**left**: up-stimulated (magenta); **middle**: down-stimulated (blue); **right**: not-stimulated (green); **b. Between condition contrasts.** Time-frequency representation (TFR) of the difference in power modulation around the trough of the SWs on which the grand-average of the up- vs. down-stimulated SWs is super-imposed (**left**), up- vs. not-stimulated SWs (**middle**), and down- vs. not-stimulated SWs (**right**). As in the main text, black lines represent the adjacent time points of the significant spatio-temporal clusters showing a difference in SW amplitude between the two trough-locked ERPs and highlighted areas in the TFR represents the adjacent time-frequency points of the significant spatio-temporal-frequency cluster showing a difference in power between conditions.*

#### SLEEP EVENT DETECTIONS

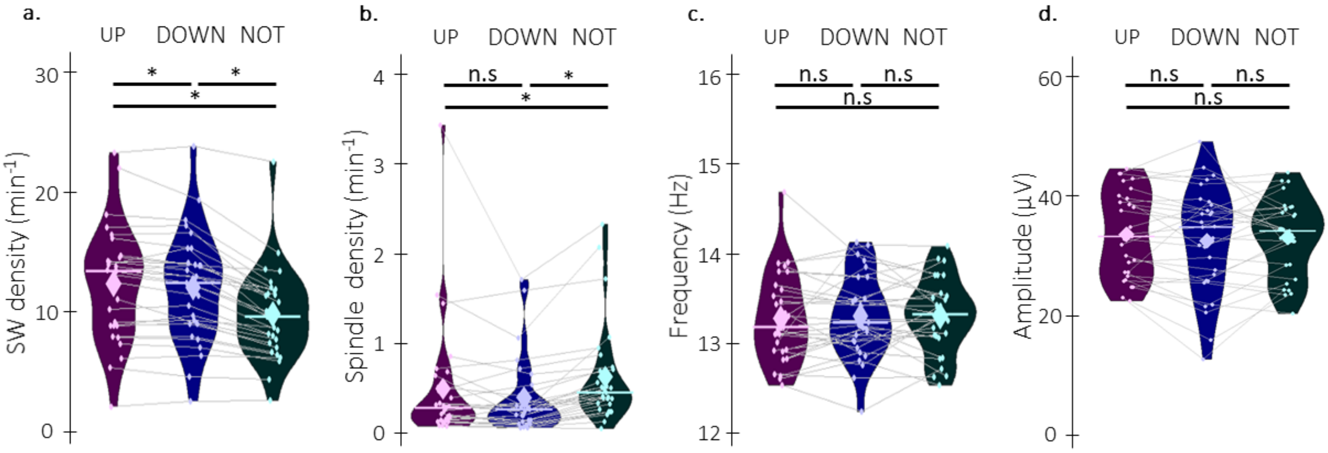

***Figure S2: Offline detected sleep events. a. SW density*** *(number of SWs per minute spent in stimulation or rest intervals) extracted from the stimulated and non-stimulated blocks on each of the 6 EEG channels and then averaged across channels was also modulated by the stimulation condition (main effect of condition: F (2,56) = 52.86, p < 0.001 (< 0.001 sphericity corrected), $\eta^2$ = 0.65). SW density was greater during both up- and down-stimulated as compared to non-stimulated blocks (up vs. not: t = 7.50, df = 29, p-value < 0.001 (< 0.001 FDR-corrected), Cohen's d = 1.37; down vs. not: t = 8.03, df = 29, p-value < 0.001 (< 0.001 FDR-corrected), Cohen's d = 1.47) and SW density was greater during up- as compared to down-stimulated blocks (up vs. down: t = -2.28, df = 29, p-value = 0.030 (0.030 FDR-corrected), Cohen's d = 0.42).* ***b. Spindle density*** *was modulated by the intervention (condition effect: F (2,56) = 10.61, p < 0.001 (< 0.001 sphericity corrected), $\eta^2$ = 0.28), such that density was lower during both up- and down-stimulated as compared to non-stimulated blocks, irrespective of the phase of the stimulation (up vs. down: t = 1.50, p-value = 0.15 (0.15 FDR-corrected), Cohen's d = 0.27; up vs. not: t = 5.88, p-value < 0.001 (< 0.001 FDR-corrected), Cohen's d = 1.07; down vs. not: t = -2.84, p-value = 0.008 (0.012 FDR-corrected), Cohen's d = 0.52).* ***c. Spindle frequency*** *did not statistically differ among stimulation conditions (F (2,56) = 0.076, p = 0.93 (0.92 sphericity corrected), $\eta^2$ = 0.0026). d.* ***Spindle amplitude*** *also did not statistically differ among stimulation conditions (F (2,56) = 0.071, p = 0.50 (0.49 sphericity corrected), $\eta^2$ = 0.0026). Violin plots: median (horizontal bar), mean (diamond), the shape of the violin plots depicts the distribution of the data. Colored points represent individual data, jittered in arbitrary distances on the x-axis within the respective violin plot to increase perceptibility.* ***n.s. = non-significant; *: p-value < 0.05***

MAIN EFFECT OF PRACTICE (ACROSS SEQUENCES) AND ITS LINEAR MODULATION BY PERFORMANCE

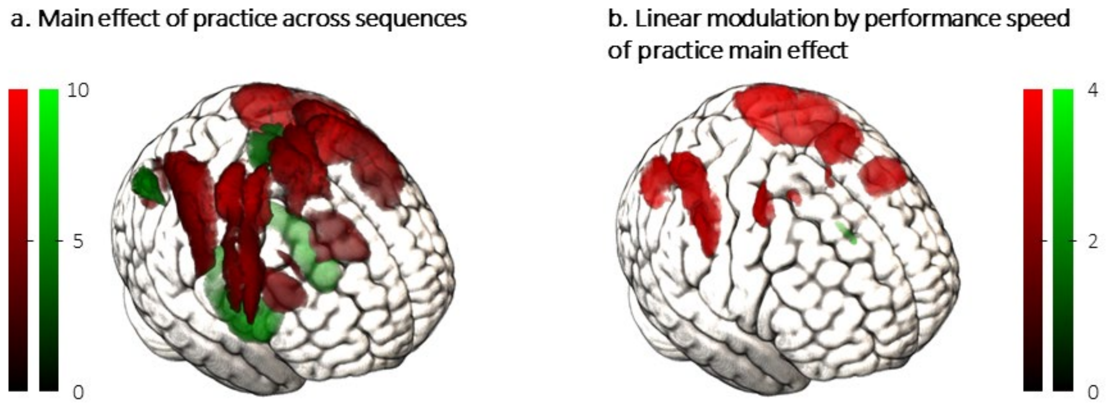

*Figure S3: Brain activity elicited by the practice of the motor sequence learning task. a. Main effect of **practice across sequences.** Task practice – as compared to rest - resulted in greater (red) activity in striatal and motor cortical regions and in less (green) activity the bilateral hippocampus and the supplementary motor area. **b. Linear modulation of brain activity as function of changes performance during task practice.** Brain activity decreased (red) with performance improvement in a network including motor and parietal regions while it increased (green) in the caudate.*

Table S1: Functional imaging results of the main effect of motor sequence learning. The significance threshold was set at $p_{uncorrected} < .005$.

| Brain Area | X mm | Y mm | Z mm | Z |
|---|---|---|---|---|
| 1. Practice vs. Rest | | | | |
| [+T] | | | | |
| L_M1 | -38 | -36 | 42 | 6.93 |
| L M1 | -40 | -10 | 62 | 6.90 |
| L SMA | -10 | -6 | 60 | 6.55 |
| R aSPL | 28 | -54 | 60 | 6.53 |
| R Putamen | 24 | -2 | 12 | 4.92 |
| L Putamen | -22 | -4 | 10 | 4.79 |
| [-T] | | | | |
| R Hippocampus | 28 | -14 | -20 | 6.29 |
| L Hippocampus | -28 | -20 | -16 | 6.17 |
| R aSPL | 46 | -72 | 42 | 5.68 |
| L aSPL | -44 | -64 | 44 | 5.28 |
| L SMA | 0 | -34 | 54 | 4.33 |
| L SMA | -12 | -32 | 60 | 3.96 |
| 2. Practice vs. Rest modulated by performance speed | | | | |

| | | | | |
|---|---|---|---|---|
| [+T] | | | | |
| R aSPL | 34 | -72 | 26 | 4.58 |
| L aSPL | -24 | -62 | 50 | 4.45 |
| L PMC | -28 | -6 | 60 | 3.71 |
| L PMC | -50 | 8 | 26 | 3.68 |
| R M1 | 30 | -6 | 52 | 3.45 |
| L SMA | -10 | 4 | 50 | 3.19 |
| [-T] | | | | |
| L Caudate | -12 | 6 | 22 | 2.98 |

FUNCTIONAL IMAGING RESULTS REPORTED IN THE MAIN TEXT

**Table S2:** Functional imaging results for the activation-based analyses. The significance threshold was set at $p_{uncorrected} < .005$.

| Brain Area | X mm | Y mm | Z mm | Z |
|---|---|---|---|---|
| **1. CHANGES IN BRAIN ACTIVITY BETWEEN SESSIONS WITHIN STIMULATION CONDITIONS** | | | | |
| **1.1. Up** | | | | |
| [Post – Pre] | | | | |
| R Putamen | 24 | 18 | -10 | 5.33 |
| R Caudate | 10 | -2 | 14 | 4.11 |
| L SMA | -6 | -12 | 56 | 5.00 |
| L M1 | -38 | -20 | 52 | 4.35 |
| R M1 | 22 | -26 | 52 | 4.28 |
| L Putamen | -24 | -8 | -2 | 4.26 |
| L Caudate | -6 | 14 | 2 | 3.80 |
| R M1 | 30 | -38 | 60 | 3.58 |
| L aSPL | -40 | -62 | 40 | 3.17 |
| L M1 | -60 | 2 | 16 | 3.03 |
| L aSPL | -56 | -26 | 46 | 3.00 |
| L aSPL | -56 | -32 | 32 | 2.97 |
| R aSPL | 58 | -18 | 28 | 2.94 |
| L M1 | -22 | -14 | 72 | 2.86 |
| [Pre – Post] No suprathreshold clusters | | | | |
| **1.2. Down** | | | | |
| [Post – Pre] | | | | |
| L SMA | -8 | -12 | 56 | 4.61 |
| L M1 | -38 | -24 | 54 | 4.52 |
| R PMC | 56 | 14 | 26 | 4.23 |
| R Putamen | 26 | -8 | -4 | 4.03 |
| R aSPL | 66 | -16 | 28 | 3.95 |
| R M1 | 30 | -38 | 60 | 3.83 |
| L Putamen | -36 | 0 | 2 | 3.60 |
| L aSPL | -60 | -30 | 34 | 3.52 |
| R Caudate | 20 | 22 | -8 | 3.29 |
| R M1 | 42 | -4 | 44 | 3.18 |

| | | | | |
|---|---|---|---|---|
| L M1 | -30 | -38 | 52 | 3.10 |
| L SMA | -16 | -14 | 60 | 2.94 |
| R Caudate | 10 | -2 | 14 | 2.89 |
| [Pre – Post] | | | | |
| R Hippocampus | 32 | -38 | -6 | 2.80 |

| **1.3. Not** | | | | |
|---|---|---|---|---|
| [Post – Pre] | | | | |
| L SMA | -2 | -4 | 46 | 4.65 |
| L Putamen | -30 | -14 | 10 | 4.29 |
| R PMC | 56 | 8 | 8 | 4.13 |
| R Putamen | 28 | -12 | -6 | 3.88 |
| L M1 | -38 | -26 | 56 | 3.71 |
| R M1 | 34 | -22 | 54 | 3.53 |
| R Caudate | 10 | 6 | 12 | 3.35 |
| R aSPL | 58 | -18 | 28 | 3.23 |
| R PMC | 50 | 0 | 52 | 3.11 |
| L aSPL | -58 | -32 | 34 | 3.10 |
| L aSPL | -42 | -62 | 44 | 3.06 |
| [Pre – Post] | | | | |
| R Hippocampus | 34 | -38 | -6 | 3.67 |
| L Hippocampus | -34 | -40 | -8 | 2.97 |

| Brain Area | X mm | Y mm | Z mm | Z |
|---|---|---|---|---|
| **2. DIFFERENCES BETWEEN CONDITIONS IN INTER-SESSION CHANGES IN BRAIN ACTIVITY** | | | | |
| **2.1. Up vs. Down** | | | | |
| [Post – Pre] | | | | |
| R Caudate | 20 | 18 | 12 | 3.41 |
| R Putamen | 22 | 4 | 12 | 3.00 |
| [Pre – Post] No suprathreshold clusters | | | | |
| **2.2. Up vs. Not** | | | | |
| [Post – Pre] | | | | |
| R Caudate | 18 | 28 | 4 | 3.12 |

| Brain Area | X mm | Y mm | Z mm | Z |
|---|---|---|---|---|
| L Caudate | -18 | 8 | 22 | 3.08 |
| R Caudate | 4 | 16 | 6 | 3.02 |
| R Hippocampus | 36 | -36 | -4 | 2.98 |
| R Caudate | 18 | 4 | 24 | 2.96 |
| [Pre – Post] No suprathreshold clusters | | | | |
| **2.3. Down vs. Not** | | | | |
| [Post – Pre] | | | | |
| R M1 | 22 | -28 | 54 | 3.42 |
| L aSPL | -36 | -46 | 34 | 3.31 |
| R Caudate | 16 | -2 | 26 | 3.22 |
| R aSPL | 36 | -42 | 34 | 3.19 |
| R SMA | 20 | -12 | 56 | 3.08 |
| [Pre – Post] No suprathreshold clusters | | | | |

| Brain Area | X mm | Y mm | Z mm | Z |
|---|---|---|---|---|
| **3. REGRESSIONS BETWEEN OVERNIGHT INCREASE IN BRAIN ACTIVITY AND TMR INDEX** | | | | |
| **3.1. Up x TMR up** | | | | |
| [Negative correlation] No suprathreshold clusters | | | | |
| [Positive correlation] | | | | |
| L Putamen | -22 | 20 | -4 | 4.08 |
| L Caudate | -18 | 24 | 10 | 4.06 |
| L Putamen | -28 | 8 | 8 | 4.05 |
| R PMC | 54 | 12 | 20 | 4.03 |
| R Putamen | 30 | 10 | 6 | 4.03 |
| R Caudate | 18 | 0 | 18 | 3.85 |
| R Caudate | 20 | 18 | 12 | 3.67 |
| R Hippocampus | 36 | -38 | -8 | 3.64 |
| R Hippocampus | 40 | -28 | -12 | 2.85 |
| L Caudate | -4 | 6 | 0 | 3.59 |
| L M1 | -24 | -30 | 52 | 3.40 |
| L M1 | -40 | -2 | 30 | 3.36 |
| R Hippocampus | 18 | -34 | 8 | 3.33 |
| R SMA | 8 | -4 | 50 | 3.30 |

| | | | | |
|---|---|---|---|---|
| R M1 | 40 | -10 | 52 | 3.23 |
| L SMA | -10 | -8 | 70 | 3.19 |
| R aSPL | 48 | -36 | 40 | 3.08 |
| R M1 | 30 | -38 | 58 | 3.05 |
| L M1 | -28 | -12 | 68 | 2.97 |
| L SMA | -20 | -12 | 54 | 2.94 |
| L Hippocampus | -30 | -44 | 0 | 2.94 |
| R M1 | 28 | -20 | 66 | 2.89 |
| R M1 | 52 | -20 | 32 | 2.87 |
| R Hippocampus | 40 | -20 | -14 | 2.85 |
| R Putamen | 34 | -8 | -6 | 2.82 |
| R M1 | 32 | -38 | 38 | 2.77 |
| **3.2. Down x TMR down** | | | | |
| [Negative correlation] No suprathreshold clusters | | | | |
| [Positive correlation] | | | | |
| R Caudate | 16 | -2 | 26 | 3.07 |
| L Hippocampus | -16 | -36 | 6 | 2.87 |
| R Hippocampus | 20 | -34 | 4 | 2.84 |
| R SMA | 4 | 2 | 58 | 2.71 |

| Brain Area | X mm | Y mm | Z mm | Z |
|---|---|---|---|---|
| **4. REGRESSIONS BETWEEN OVER NIGHT CHANGES IN BRAIN ACTIVITY AND EEG** | | | | |
| **4.1. Up x sigma power up** | | | | |
| [Negative correlation] | | | | |
| R PMC | 56 | 8 | 10 | 2.94 |
| L M1 | -50 | -8 | 38 | 2.87 |
| [Positive correlation] No suprathreshold clusters | | | | |
| **4.2. Down x sigma power down** | | | | |
| [Negative correlation] No suprathreshold clusters | | | | |
| [Positive correlation] | | | | |
| L Caudate | -10 | 18 | -6 | 2.72 |
| **4.3. Up x peak amplitude up** | | | | |

| | | | | |
|---|---|---|---|---|
| [Negative correlation] No suprathreshold clusters | | | | |
| [Positive correlation] | | | | |
| R Pallidum | 18 | -4 | -4 | 3.49 |
| R aSPL | 56 | -42 | 54 | 2.85 |
| L M1 | -46 | -16 | 48 | 2.72 |

| 4.4. Down x peak amplitude down | | | | |
|---|---|---|---|---|
| [Negative correlation] No suprathreshold clusters | | | | |
| [Positive correlation] | | | | |
| L M1 | -24 | -20 | 68 | 3.34 |
| L M1 | -44 | -16 | 52 | 3.33 |
| L aSPL | -62 | -20 | 40 | 3.09 |
| L M1 | -46 | -10 | 26 | 3.02 |
| L M1 | -20 | -16 | 70 | 2.97 |
| R M1 | 48 | -8 | 26 | 2.86 |
| R M1 | 44 | -14 | 52 | 2.78 |

Table S3: Functional imaging results for the connectivity analyses. The significance threshold was set at $p_{uncorrected} < .005$.

| Brain Area | X mm | Y mm | Z mm | Z |
|---|---|---|---|---|
| 1. OVERNIGHT CHANGES IN CONNECTIVITY WITHIN CONDITIONS | | | | |
| 1.1. RIGHT CAUDATE | | | | |
| 1.1.1. Up | | | | |
| [Post – Pre] No suprathreshold clusters | | | | |
| [Pre – Post] No suprathreshold clusters | | | | |
| 1.1.2. Down | | | | |
| [Post – Pre] No suprathreshold clusters | | | | |
| [Pre – Post] No suprathreshold clusters | | | | |
| 1.1.3. Not | | | | |
| [Post – Pre] | | | | |
| L Hippocampus | -32 | -20 | -14 | 2.70 |
| [Pre – Post] No suprathreshold clusters | | | | |
| 1.2. RIGHT PUTAMEN | | | | |
| 1.2.1. Up | | | | |
| [Post – Pre] No suprathreshold clusters | | | | |
| [Pre – Post] | | | | |
| R Putamen | 30 | 12 | -2 | 2.81 |
| 1.2.2. Down | | | | |
| [Post – Pre] | | | | |
| L M1 | -46 | -8 | 26 | 3.03 |
| R M1 | 28 | -8 | 44 | 2.92 |
| L M1 | -52 | 2 | 48 | 2.71 |
| L M1 | -32 | -18 | 48 | 2.69 |
| L SMA | -22 | -12 | 58 | 2.67 |
| [Pre – Post] No suprathreshold clusters | | | | |

| | | | | |
|---|---|---|---|---|
| **1.2.3. Not** | | | | |
| [Post – Pre] | | | | |
| R Ventral striatum | 14 | 18 | -10 | 3.02 |
| [Pre – Post] No suprathreshold clusters | | | | |

| | | | | |
|---|---|---|---|---|
| **1.3. RIGHT HIPPOCAMPUS** | | | | |
| **1.3.1. Up** | | | | |
| [Post – Pre] No suprathreshold clusters | | | | |
| [Pre – Post] | | | | |
| R PMC | 40 | 4 | 26 | 3.15 |
| R PMC | 42 | -4 | 56 | 3.08 |
| R aSPL | 36 | -66 | 28 | 3.00 |
| L M1 | -38 | -4 | 56 | 2.74 |
| R aSPL | 32 | -52 | 52 | 2.72 |

| | | | | |
|---|---|---|---|---|
| **1.3.2. Down** | | | | |
| [Post – Pre] No suprathreshold clusters | | | | |
| [Pre – Post] | | | | |
| L M1 | -60 | -4 | 22 | 2.73 |

| |
|---|
| **1.3.3. Not** |
| [Post – Pre] No suprathreshold clusters |
| [Pre – Post] No suprathreshold clusters |

| Brain Area | X mm | Y mm | Z mm | Z |
|---|---|---|---|---|
| **2. OVERNIGHT CHANGES IN CONNECTIVITY BETWEEN CONDITIONS** | | | | |
| **2.1. RIGHT CAUDATE** | | | | |
| **2.1.1. Up vs. Down** | | | | |
| [Post – Pre] No suprathreshold clusters | | | | |
| [Pre – Post] | | | | |
| R aSPL | 54 | -36 | 56 | 2.80 |

| |
|---|
| **2.1.2. Up vs. not** |
| [Post – Pre] No suprathreshold clusters |

| | | | | |
|---|---|---|---|---|
| [Pre – Post] | | | | |
| L aSPL | -50 | -50 | 48 | 2.97 |
| R aSPL | 50 | -36 | 56 | 2.77 |
| **2.1.3. Down vs. Not** | | | | |
| [Post – Pre] No suprathreshold clusters | | | | |
| [Pre – Post] | | | | |
| L Pallidum | -22 | -14 | -4 | 2.81 |
| **2.2. RIGHT PUTAMEN** | | | | |
| **2.2.1. Up vs. Down** | | | | |
| [Post – Pre] No suprathreshold clusters | | | | |
| [Pre – Post] | | | | |
| R aSPL | 28 | -46 | 42 | 3.60 |
| L M1 | -22 | -14 | 52 | 3.17 |
| L M1 | -32 | -20 | 50 | 3.15 |
| R Putamen | 26 | 12 | -4 | 2.93 |
| L SMA | -10 | -10 | 54 | 2.87 |
| **2.2.2. Up vs. not** | | | | |
| [Post – Pre] No suprathreshold clusters | | | | |
| [Pre – Post] No suprathreshold clusters | | | | |
| **2.2.3. Down vs. not** | | | | |
| [Post – Pre] No suprathreshold clusters | | | | |
| [Pre – Post] No suprathreshold clusters | | | | |
| **2.3. RIGHT HIPPOCAMPUS** | | | | |
| **2.3.1. Up vs. Down** | | | | |
| [Post – Pre] No suprathreshold clusters | | | | |
| [Pre – Post] No suprathreshold clusters | | | | |
| **2.3.2. Up vs. not** | | | | |
| [Post – Pre] No suprathreshold clusters | | | | |
| [Pre – Post] | | | | |

| | | | | |
|---|---|---|---|---|
| L M1 | -26 | -22 | 54 | 3.46 |
| R SMA | 12 | -12 | 48 | 3.36 |
| R M1 | 60 | -24 | 48 | 3.19 |
| L M1 | -26 | -14 | 54 | 3.05 |
| R SMA | 22 | -8 | 52 | 3.02 |
| L aSPL | -40 | -42 | 48 | 2.96 |
| R M1 | 28 | -38 | 54 | 2.91 |
| R M1 | 48 | -10 | 52 | 2.80 |
| R M1 | 24 | -28 | 54 | 2.79 |
| R SMA | 14 | -4 | 68 | 2.69 |
| L aSPL | -26 | -52 | 54 | 2.69 |
| **2.3.3. Down vs. not** | | | | |
| [Post – Pre] No suprathreshold clusters | | | | |
| [Pre – Post] | | | | |
| L SMA | 0 | -34 | 54 | 3.59 |
| R M1 | 22 | -14 | 56 | 3.21 |
| L aSPL | -30 | -46 | 44 | 3.12 |
| L M1 | -54 | -10 | 44 | 3.10 |
| L M1 | -48 | -10 | 26 | 3.05 |
| R M1 | 48 | -10 | 34 | 3.00 |
| R M1 | 54 | -22 | 46 | 2.85 |
| Caudate_L | -18 | -14 | 20 | 2.74 |
| R SMA | 12 | -12 | 46 | 2.72 |
| R M1 | 62 | -2 | 36 | 2.69 |

| Brain Area | X mm | Y mm | Z mm | Z |
|---|---|---|---|---|
| **3. REGRESSION BETWEEN OVERNIGHT CHANGES IN CONNECTIVITY WITHIN CONDITION AND TMR INDEX** | | | | |
| 3.1. RIGHT CAUDATE | | | | |
| **3.1.1. Overnight up x TMR up** | | | | |
| [Negative correlation] | | | | |
| R aSPL | 44 | -46 | 50 | 2.84 |
| [Positive correlation] No suprathreshold clusters | | | | |

##### 3.1.2. Overnight down x TMR down

| | | | | |
|---|---|---|---|---|
| [Negative correlation] No suprathreshold clusters | | | | |
| [Positive correlation] | | | | |
| L Hippocampus | -16 | -40 | 6 | 3.30 |
| R Caudate | 16 | -2 | 26 | 3.24 |
| R Ventral striatum | 4 | 6 | 0 | 2.85 |
| R Hippocampus | 18 | -38 | 6 | 2.72 |

#### 3.2. RIGHT PUTAMEN

##### 3.2.1. Overnight up x TMR up

| | | | | |
|---|---|---|---|---|
| [Negative correlation] | | | | |
| R aSPL | 44 | -42 | 44 | 2.78 |
| R aSPL | 38 | -60 | 42 | 2.65 |
| [Positive correlation] | | | | |
| R Pallidum | 22 | -8 | -6 | 2.95 |
| L Caudate | -16 | 6 | 22 | 2.86 |

##### 3.2.2. Overnight down x TMR down

| | | | |
|---|---|---|---|
| [Negative correlation] | | | |
| R Putamen | 28 | -2 | 14 |
| R M1 | 26 | -8 | 44 |
| R Caudate | 12 | -14 | 20 |
| [Positive negative correlation] No suprathreshold clusters | | | |

#### 3.3. RIGHT HIPPOCAMPUS

##### 3.3.1. Overnight up x TMR up

| | | | | |
|---|---|---|---|---|
| [Negative correlation] | | | | |
| R aSPL | 40 | -70 | 34 | 3.71 |
| R Putamen | 32 | -8 | 6 | 3.10 |
| R SMA | 8 | -20 | 62 | 2.82 |
| [Positive correlation] | | | | |
| R Hippocampus | 28 | -36 | -4 | 2.74 |

| Brain Area | x | y | z | Z |
|---|---|---|---|---|
| **4. REGRESSION BETWEEN OVERNIGHT CHANGES IN CONNECTIVITY WITHIN CONDITION AND EEG** | | | | |
| **4.1. RIGHT CAUDATE** | | | | |
| **4.1.1. Up x Sigma power up** | | | | |
| [Negative correlation] | | | | |
| R aSPL | 44 | -40 | 38 | 3.41 |
| R aSPL | 36 | -68 | 42 | 2.70 |
| [Positive correlation] | | | | |
| L SMA | -10 | -22 | 58 | 3.41 |
| R Caudate | 14 | 24 | 10 | 3.36 |
| R Hippocampus | 38 | -14 | -22 | 3.33 |
| R Ventral striatum | 6 | 8 | -6 | 3.30 |
| R Putamen | 24 | 8 | -12 | 2.80 |
| **4.1.2. Down x Sigma power down** | | | | |
| [Negative correlation] | | | | |
| L Hippocampus | -24 | -14 | -8 | 3.68 |
| R Ventral striatum | 12 | 6 | -8 | 3.65 |
| R Hippocampus | 20 | -18 | -24 | 3.55 |
| L Ventral striatum | -8 | 8 | -12 | 3.49 |
| L aSPL | -38 | -70 | 44 | 3.41 |
| R aSPL | 40 | -72 | 48 | 3.26 |
| L Putamen | -34 | 4 | -8 | 3.20 |
| [Positive correlation] No suprathreshold clusters | | | | |
| **4.2. RIGHT PUTAMEN** | | | | |
| **4.2.1. Up x sigma power up** | | | | |
| [Negative correlation] No suprathreshold clusters | | | | |
| [Positive correlation] | | | | |
| L Caudate | -18 | 24 | 10 | 3.13 |
| **4.2.2. Down x sigma power down** | | | | |
| [Negative correlation] | | | | |
| L Hippocampus | -18 | -10 | -8 | 3.25 |

| |
|---|
| [Positive correlation] No suprathreshold clusters |

#### 4.3. RIGHT HIPPOCAMPUS

##### 4.3.1. Up x sigma power up

| | | | | |
|---|---|---|---|---|
| [Negative correlation] | | | | |
| R M1 | 26 | -4 | 48 | 3.15 |
| [Positive correlation] No suprathreshold clusters | | | | |

##### 4.3.2. Down x sigma power down

[Negative correlation] No suprathreshold clusters

[Positive correlation] No suprathreshold clusters

#### 4.4. RIGHT CAUDATE

##### 4.4.1. Up x peak amplitude up

[Negative correlation] No suprathreshold clusters

[Positive correlation] No suprathreshold clusters

##### 4.4.2. Down x peak amplitude down

| | | | | |
|---|---|---|---|---|
| [Negative correlation] | | | | |
| R Hippocampus | 14 | -30 | -10 | 4.62 |
| L Hippocampus | -16 | -8 | -16 | 4.26 |
| R PMC | 50 | -2 | 54 | 4.25 |
| R SMA | 6 | 2 | 64 | 4.24 |
| L M1 | -62 | -20 | 28 | 3.90 |
| L aSPL | -44 | -44 | 38 | 3.84 |
| L M1 | -62 | 4 | 16 | 3.67 |
| L M1 | -34 | -4 | 42 | 3.36 |
| L Caudate | -6 | 2 | 0 | 3.54 |
| R M1 | 62 | -2 | 14 | 3.35 |
| L M1 | -24 | -28 | 58 | 3.32 |
| R aSPL | 66 | -16 | 28 | 3.31 |
| R Caudate | 12 | -8 | 18 | 3.26 |
| R M1 | 26 | -46 | 46 | 3.18 |
| R PMC | 56 | 8 | 8 | 3.14 |
| L M1 | -44 | -12 | 44 | 2.99 |

| | | | | |
|---|---|---|---|---|
| L Caudate | -14 | 20 | 6 | 2.89 |
| R Ventral striatum | 2 | 8 | -2 | 2.86 |
| L Putamen | -26 | -18 | 10 | 2.82 |
| R Caudate | 8 | 14 | 8 | 2.72 |
| [Positive correlation] No suprathreshold clusters | | | | |

#### 4.5. RIGHT PUTAMEN

##### 4.5.1. Up x peak amplitude up

[Negative correlation] No suprathreshold clusters

[Positive correlation] No suprathreshold clusters

##### 4.5.2. Down x peak amplitude down

| | | | | |
|---|---|---|---|---|
| [Negative correlation] | | | | |
| L aSPL | -36 | -42 | 36 | 4.56 |
| | *-38* | *-42* | *36* | |
| R aSPL | 34 | -42 | 48 | 3.93 |
| R Hippocampus | 32 | -22 | -12 | 3.74 |
| R aSPL | 30 | -66 | 38 | 3.56 |
| R SMA | 12 | -2 | 66 | 3.55 |
| R M1 | 58 | 12 | 36 | 3.54 |
| R PMC | 28 | -12 | 60 | 3.16 |
| L M1 | -48 | 4 | 32 | 3.15 |
| R M1 | 40 | 0 | 46 | 3.06 |
| R Hippocampus | 26 | -38 | -6 | 3.01 |
| R SMA | 8 | 4 | 48 | 2.98 |
| R aSPL | 34 | -66 | 58 | 2.94 |
| L Pallidum | -16 | -2 | -2 | 2.92 |
| R Hippocampus | 30 | -4 | -28 | 2.86 |
| L Putamen | -26 | -14 | 12 | 2.79 |
| L M1 | -28 | -14 | 62 | 2.75 |
| L Hippocampus | -26 | -36 | -10 | 2.70 |
| [Positive correlation] No suprathreshold clusters | | | | |

#### 4.6. RIGHT HIPPOCAMPUS

##### 4.6.1. Up x peak amplitude up

[Negative correlation] No suprathreshold clusters

| |
|---|
| [Positive correlation] No suprathreshold clusters |

##### 4.6.2. Down x peak amplitude down

| | | | | |
|---|---|---|---|---|
| [Negative correlation] | | | | |
| L SMA | -6 | -28 | 56 | 3.42 |
| L PMC | -50 | 8 | 16 | 3.17 |
| L M1 | -32 | -18 | 42 | 3.17 |
| L aSPL | -46 | -56 | 46 | 2.80 |
| R Caudate | 8 | 16 | 12 | 3.05 |
| L Caudate | -8 | 6 | 14 | 3.03 |
| L Caudate | -12 | 14 | 6 | 2.94 |
| L M1 | -24 | -28 | 58 | 2.98 |
| R SMA | 18 | -26 | 58 | 2.95 |
| R M1 | 56 | -2 | 18 | 2.71 |
| [Positive correlation] No suprathreshold clusters | | | | |

### CONNECTIVITY REGRESSIONS SUPPLEMENTARY RESULTS

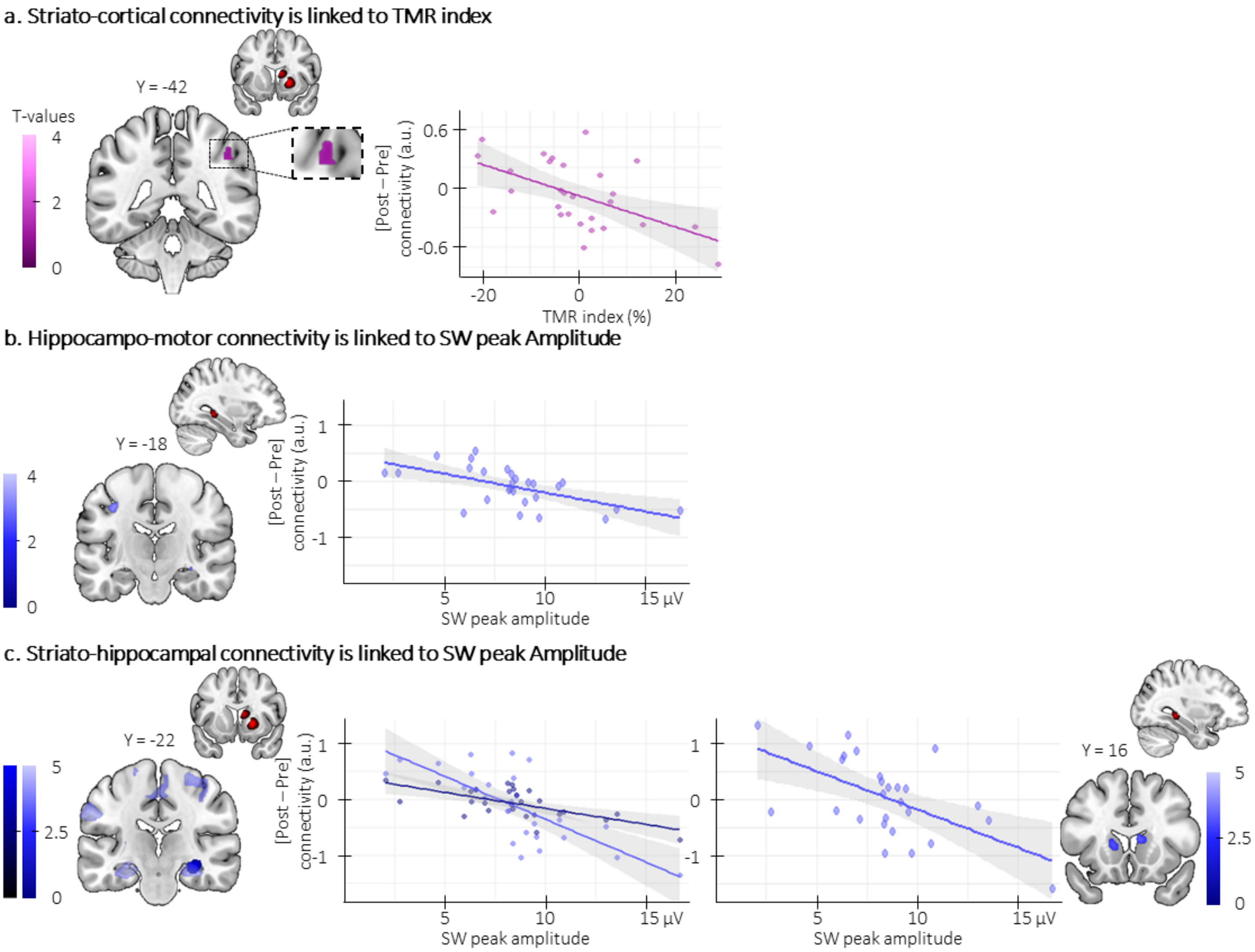

*Figure S4: Regressions between connectivity, behavior and EEG metrics. A Striato-cortical connectivity and TMR inde.* *The overnight decrease in striato-motor connectivity was linked to the TMR index for the up-reactivated sequence (putamen-right aSPL: x = 44, y = -42 z = 44, $p_{SVC}$ = 0.053) such that the greater the decrease in connectivity (y-axis), the greater the performance improvement on the up-reactivated as compared to the not-reactivated sequence (x-axis).* ***b. Hippocampo-motor connectivity and SW peak amplitude.*** *The overnight decrease in hippocampo-motor connectivity was linked to the SW peak amplitude in the down-stimulated condition (hippocampus-left M1: x = -32, y = -18, z = 42, $p_{SVC}$ = 0.02) such that the greater the decrease in connectivity (y-axis), the greater the SW peak amplitude during the night(x-axis).* ***c. Striato-hippocampal connectivity and SW peak amplitude.*** *The overnight decrease in striato-hippocampal connectivity was linked to the SW peak amplitude in the down-stimulated condition (**left panel**, Caudate-right hippocampus (pale blue): x = 14, y = -32, z = -8, $p_{SVC}$ < 0.001; putamen-right hippocampus (dark blue): x = 32, y = -22, z = -12, $p_{SVC}$ = 0.004; **right panel**, hippocampus-right caudate: x = 8, y = 16, z =12, pSVC = 0.027) such that the greater the decrease in connectivity (y-axis), the greater the SW peak amplitude during the night (x-axis). Activations maps are displayed on a T1-weighted template image with a threshold of p < 0.005 uncorrected. a.u.: arbitrary units. Violin plots: median (horizontal bar), mean (diamond), the shape of the violin plots depicts the distribution of the data.*

**Tables S4**. List of the deviations from the pre-registered analyses followed by their justification. These deviations are marked with a # in the main manuscript.

| | Pre-registered | Final report |
|---|---|---|
| 1 | Task-based functional volumes of each participant will be realigned to the first image of each session and then realigned to the mean functional image computed across MRI runs of the respective participant using rigid body transformations **and corrected for inhomogeneities in the magnetic field (i.e., unwarped) using the field map images**. | Additionally, to correct for distortions due to magnetic field inhomogeneities, field maps (TR = 1500 ms; TE = 3.5 ms; flip angle = 90°; 42 transverse slices; slice thickness = 3.75 mm; interslice gap = 0 mm; voxel size = 3.75 × 3.75 × 3.75 mm3; field of view = 240 × 240 × 157.5 mm3; matrix = 64 × 64) were collected […]. **However, these sequences were not included in the final analysis pipeline.** |
| | **Justification**: The introduction of the field maps in the processing pipeline induced unforeseen image distortions in some participants. We therefore elected to not use the field maps in the final analysis pipeline in order to keep the preprocessing consistent across individuals. | |
| 2 | Individual trials (i.e., reaction times to visual stimuli during the MSL tasks) will be excluded from further analysis if the trial deviates more than three standard deviations from the block's average reaction time. | Correct trials were excluded from the analyses if they were outlier trials based on **John Tukey's method of leveraging the Interquartile**. |
| | **Justification**: The John Tukey's method of leveraging the Interquartile was used for outlier exclusion as the median was the statistical average used in this study. | |
| 3 | Only correct stimulation (i.e. auditory cues sent during true SW as defined in [1] and refined later in [2]) will be considered for the analyses. To analyze oscillatory activity **evoked by the auditory stimulations**, we will compute Time-Frequency representations (TFR) averaged across all trials of each condition. Individual TFRs will be converted into baseline relative change of power (baseline from -300 ms to -100 ms relative to cue onset), thus highlighting power modulation following the auditory cues. | Event-related data analyses included **trough-locked potentials and oscillatory activity** and were performed with down sampled data (100Hz). Trough-locked responses were obtained by segmenting the data into epochs time-locked to the trough of the detected SW (from -2 to 2 sec) separately for the stimulated-up, stimulated-down and not-stimulated SW and averaged across all trials.<br>[…]<br>Individual TFRs were converted into change of power relative **to the entire period around the SW trough (from -2s to 2s relative to trough).** |
| | **Justification**: Instead of the auditory cue-locked analysis, we chose to perform trough-locked analyses to reduce the influence of the phase of the SW on activity evoked by the stimulation. Specifically, as auditory cues are sent at different SW phases, the corresponding auditory-evoked activity is embedded into high signal amplitude fluctuation, which prevents us from directly comparing conditions. This deviation also allowed us to substitute the SW amplitude analysis performed on the discrete sleep events with more comprehensive (i.e., across all time points in the analysis window) and conservative trough-locked potential analysis using the same approach as the pre-registered oscillatory activity analysis, i.e., using Cluster-based permutation approach. Nonetheless, the corresponding hypotheses remained unchanged. Namely, (i) up-stimulated SW amplitude will be boosted as compared to down- and not-stimulated SW whereas down-stimulated SW amplitude will be impaired as compared to up- and not-stimulated SWs. (ii) CL-TMR will modulate the power in the sigma frequency band regardless of the stimulation phase. Finally, as the analyses were focused on physiological events and not stimulus-evoked responses for which the time preceding the cue is supposed to represent a brain activity-free moment, we choose to use the entire period around the SW to perform a normalization of the TF power similar as in R. | |
| 4 | The power in the sigma band will be compared between the reactivated-up and the control SWs and between the reactivated-down and the control SWs separately using cluster-based permutation tests (Maris & Oostenveld, 2007) using an alpha threshold at 0.025 (Bonferonni corrected for two comparisons). | To identify significant evoked power modulation, TFR locked to the SW trough were compared 2-by-2 using a CBP tests between 5 to 30 Hz and from -1.5 to 1.5 sec relative to cue onset to exclude border effects and **Bonferonni corrected for three multiple comparisons**. |

| | | |
|---|---|---|
| | **Justification**: For completeness, we decided to also compare the two stimulated conditions. Therefore, the number of tests was increased to 3 and the Bonferroni correction needed to be adjusted. | |
| 5 | *SW-spindle coupling*: Finally, we aim to determine whether the amplitude of the oscillatory signal is coupled to the phase of the delta band using the tensorPac[4] open-source Python toolbox. The event-related PAC will be computed in relation to the trough of the SWs. | Analysis not performed. |
| | **Justification**: As per our new analysis that focused on trough-locked oscillatory activity modulation, the phase-amplitude coupling analysis became redundant. | |
| 6 | Linear contrasts will be generated at the individual level to test for (1) the main effect of practice (across sequences) and its linear modulation by performance, (2) the main effect of practice for each sequence (reactivated-up, reactivated-down, and non-reactivated) and their linear modulation by performance and (3) difference in brain responses between sequences practiced (reactivated-up vs. reactivated-down vs. non-reactivated) and their linear modulation by performance. These contrasts will be written within each task run (pre-night training, **pre-night test**, post-night training) as well as between task runs. | Pre-night test linear contrasts are not reported in the present study. |
| | **Justification**: The imaging data acquired during the pre-night test run were included in the model outlined in the pre-registration. However, the corresponding contrasts were not considered of interest as the number of volumes acquired during the three blocks of practice (for each sequence) was lower than expected (participant reached a high level of performance at the end of training which reduced time on task / number of volumes acquired for this session). We optimized the number of blocks in this pre- night test run in order to provide a reflection of end-of-training behavioral performance after allowing for the dissipation of fatigue effects (see [5]), which made these runs less suitable for imaging analyses, especially when performance reaches high levels. | |

PARTICIPANT CHARACTERISTICS

**Table S5**. Participants' characteristics, sleep and vigilance. Group averaged (+/- 95% confidence interval) sleep characteristics leading up to the experimental session and vigilance assessments at time of testing (N=31 if not specified)

| | | | | |
|---|---|---|---|---|
| **Number of datasets per analysis** | | | | |
| Behavior & fMRI | | 28 | | |
| EEG and polysomnography | | 30 | | |
| Closed-loop stimulation | | 31 | | |
| Complete datasets | | 27 | | |
| **Participant characteristics** | | | | |
| Age (yrs) | | 23.7 ranging from 18 to 30 | | |
| Edinburgh Handedness [6] | | 89.0 [84.1 - 95.9] | | |
| Epworth Sleepiness Scale [7] | | 6.0 [4.8 – 7.2] | | |
| Beck Depression Scale [8] | | 3.1 [1.9 - 4.3] | | |
| Beck Anxiety Scale [9] | | 2.6 [1.6 – 3.7] | | |
| PSQI [10] | | 2.6 [2.1 - 3.1] | | |
| Chronoscore (CRQ)[11] | | 53.1 [50.3 - 55.1] | | |
| **Sleep duration** [a] (N = 30) | | | | |
| Night 1 (minutes) | | 485.5 [471.3 - 499.6] | | |
| Night 2 (minutes) | | 492.9 [477.9 - 507.9] | | |
| Night 3 (minutes) | | 499.7 [486.6 - 512.8] | | |
| **St. Mary's questionnaire** | | *Quality* | *Duration (minutes)* | |
| Night 3 | | 4.1 [3.9 - 4.4] | 480.3 [463.2 - 497.4] | |
| Experimental Night | | 3.7 [3.4 – 4.0] | 435.4 [422.8 – 448.0] | |
| One-way rmANOVA results | | F(1,30) = 5.18; p = 0.03 | F(1,30) = 19.09; p < 0.001 | |
| **Psychomotor Vigilance Task**[b] (N = 26) | | | | |
| Pre-night RT (ms) | | 298.1 [293.9 – 302.3] | | |
| Post-night RT (ms) | | 302.4 [298.4 – 306.3] | | |
| One-way rmANOVA results | | F(1,25) = 0.7; p = 0.42 | | |
| **Stanford sleepiness score** | | | | |
| Pre-night Session | | 2.2 [1.9 - 2.4] | | |
| Post-night Session | | 2.2 [1.9 - 2.5] | | |
| One-way rmANOVA results | | F(1,30) = 0.18; p = 0.68 | | |
| **Sleep characteristics** (N = 30) | | | | |
| Time allowed to sleep (minutes) | | 450 [436.8 – 463.1] | | |
| Total Sleep Time[c] (minutes) | | 374.5 [354.9 - 354.9] | | |
| Sleep Efficiency[d] (percentage) | | 83.3 [79.6 - 87] | | |
| NREM1 Latency (minutes) | | 13.7 [17.7 - 9.7] | | |
| Time awake (minutes) | | 39.7 [25.9 - 53.5] | | |
| Time in NREM1 Sleep (minutes) | | 34.9 [28.9 - 41.0] | | |
| Time in NREM2 Sleep (minutes) | | 229.4 [216.6 - 242.1] | | |
| Time in NREM3 Sleep (minutes) | | 74.5 [63.3 - 85.7] | | |
| Time in REM Sleep (minutes) | | 70.6 [61.8 - 79.4] | | |
| **Participants reaching** (N = 30) | | | | |
| NREM3 sleep | | 30 | | |
| REM sleep | | 30 | | |
| **Algorithm accuracy** [e] (N = 31) | | | | |
| True positive | Number | 611.7 [527.6 – 695.9] | 622.9 [495.2 – 750.7] | 600.5 [483.3 - 717.7] |
| | Percentage | 82.2 % [79.6 – 84.8] | 89.5 % [87.8 - 91.2] | 74.9 % [71.7 - 78.1] |
| **Number of Auditory cues** (N = 30) | | *All cues* | *Up-reactivated cues* | *Down-reactivated cues* |
| During all stages | | 1492.2 [1228.0 -1756.5] | 700.6 [566.0 – 835.1] | 791.7 [660.1 – 923.2] |
| During wake | | 8.3 [4.6 - 11.9] | 2.7 [1.5 – 4.0] | 5.5 [2.1 - 8.9] |
| During NREM1 Sleep | | 6.3 [3.2 - 9.3] | 2.5 [0.9 - 4.1] | 3.7 [1.7 - 5.8] |
| During NREM2-3 Sleep | | 1476.5 [1211.2 - 1741.8] | 694.9 [560.0 - 829.9] | 781.6 [649.5 - 913.7] |
| During REM Sleep | | 1.2 [0.5 – 1.9] | 0.4 [0.1 - 0.7] | 0.8 [0.3 - 1.4] |
| Accuracy [f] | | 98.5 [97.6 - 99.3] | 98.8 [98.1 - 99.4] | 98.2 [97.2 - 99.3] |

*Notes*. Values are means [lower and upper limit of the 95% Confidence Interval - CI]. PSQI = Pittsburgh Sleep Quality Index; CRQ = Circadian Rhythm Questionnaire. REM: Rapid Eye Movement. [a] Sleep duration was computed as the mean across the actigraphy data and the sleep diary for the three nights before the experimental day (except for three participants for which only the sleep diary data were available, both actigraphy data and sleep diary were missing for one participant). [b] Median of reaction times computed across the 100 trials of each session. REM: Rapid Eye Movement. [c] Total sleep time was computed as the total time spent in stages NREM2-3 and REM sleep. [d] Sleep efficiency was computed as the percent of time asleep (namely in NREM2-3 and in REM sleep) relative to the total time in bed (specifically, from lights off to lights on). [e] Algorithm accuracy was based on Fpz which was recorded for all 31 participants. [f] Percentage of auditory cues correctly sent during NREM2 and NREM3 sleep based on the PSG scoring available for 30 participants.

CONFIRMATORY ANALYSIS ON ACCURACY

A one-way repeated measure analysis of variance (rmANOVA) performed on offline changes in performance accuracy with Condition (reactivated-up vs. reactivated-down vs. non-reactivated) as within-subject factor showed no significant Condition effect (F (2,54) = 0.12, p = 0.89 (0.87 sphericity corrected), $\eta^2$ = 0.0043; Figure S5b).

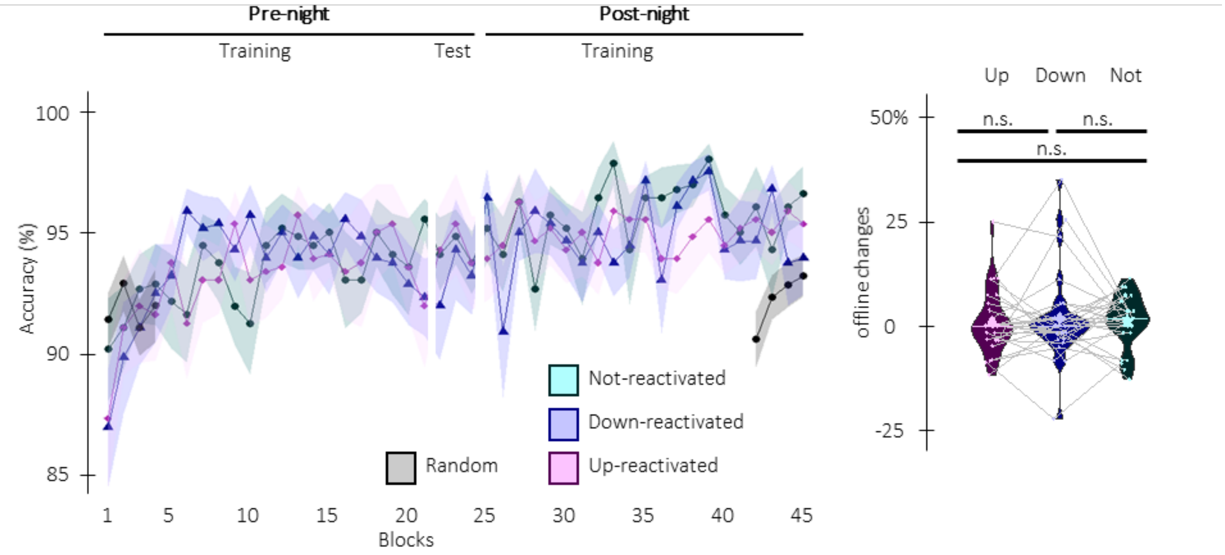

*Figure S5: Behavioral results. a. Performance accuracy (mean percentage of* correct responses*) across participants plotted as a function of blocks of practice during the pre and post-night sessions (+/- standard error) for the up-reactivated (magenta), the down-reactivated (blue) and the not-reactivated (green) sequences. See negative control analyses below for results of the statistical analyses. b. Offline changes in performance accuracy (% change between the average of the three blocks of pre-night test and the first three blocks of post-night training) averaged across participants. There was no main effect pf condition. Violin plots: median (horizontal bar), mean (diamond), the shape of the violin plots depicts the distribution of the data. Colored points represent individual data, jittered in arbitrary distances on the x-axis within the respective violin plot to increase perceptibility. For each individual, performance on the different conditions are connected with a line between violin plots. n.s: non-significant.*

NEGATIVE CONTROL ANALYSES

*Equivalent vigilance during each behavioral session*

As presented in supplementary Table S5, the one-way rmANOVAs on both the median RTs of the PVT and the Stanford Sleepiness Scale score with Session as two-level factor (pre-night and post-night) showed no significant main effect of session (PVT: $F(1,25) = 0.7$; $p = 0.42$; SSS: $F(1,30) = 0.18$; $p = 0.68$), suggesting that vigilance did not differ between the two testing sessions.

*Equivalent baseline performance between the three movement sequences*

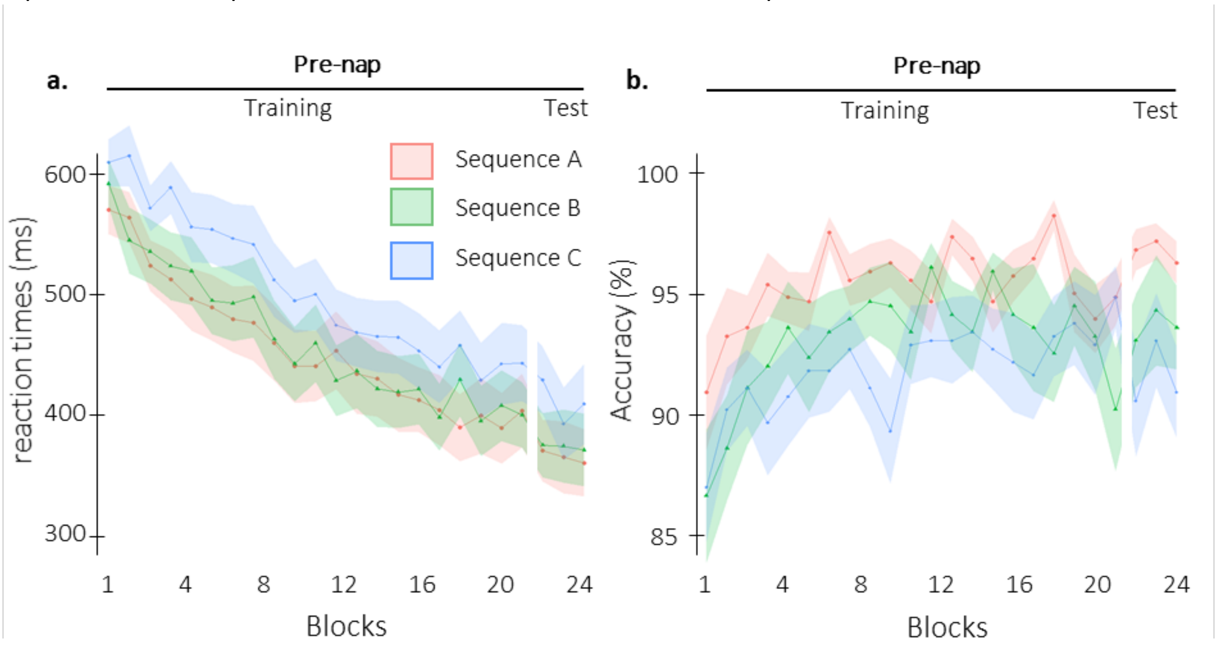

*Figure S6: Behavioral results per sequence. a. Performance speed (mean reaction time in ms +/- standard error in shaded regions) across participants plotted as a function of blocks of practice during the pre-night session (Training and Test) for the sequence A (4 7 2 8 3; red), the sequence B (1 6 3 5 2; green), and the sequence C (7 3 8 4 6; blue). b. Performance accuracy (mean percentage of* correct responses *+/- standard error in shaded regions) across participants plotted as a function of blocks of practice during the pre-night session (Training and Test) for sequences A, B and C.*

We assessed whether performance (speed and accuracy) during initial training significantly differed between the three movement sequences irrespective of the reactivation condition. To do so, two two-way rmANOVAs with sequence (A vs. B vs. C) and Block (1$^{st}$ rmANOVA on the 21 blocks of the pre-night training and 2$^{nd}$ rmANOVA on the 3 blocks of the pre-night test) as within-subject factors were performed on the sequential SSRT performance.

Analyses of the pre-night training data indicated differences in movement speed among the 3 sequences (main effect of Sequence ($F(2,54) = 20.72$, p-value $< 0.001$, $\eta^2 = 0.43$, Figure S6a), a result that can be attributed to slower RTs for Sequence C. However, performance improved similarly among sequences (main effect of Block: $F(20,540) = 42.5$, p-value $< 0.001$, $\eta^2 = 0.61$; Block by Sequence interaction: $F(40,1080)= 1.19$, p-value $= 0.20$, $\eta^2 = 0.04$). In terms of accuracy, results showed a similar pattern with a main effect of Sequence ($F(2,54) = 5.9$, p-value $= 0.0048$, $\eta^2 = 0.18$; Figure S6b) ), a result that can be attributed to better performance for Sequence C. However, accuracy improved similarly among sequences ($F(20,540) = 3.0$, p-value $< 0.001$, $\eta^2 = 0.10$; Block by Sequence interaction ($F(40,1080)= 1.2$, p-value $= 0.18$, $\eta^2 = 0.04$).

Analyses of the pre-night test data showed that sequence C was still performed significantly slower than the others sequence (main effect of Sequence ($F(2,54) = 20.72$, p-value $< 0.001$ ($< 0.001$ sphericity corrected), $\eta^2 = 0.30$, Figure S6a), yet performance remained similarly stable across blocks among sequences (main effect of Block: $F(20,54) = 1.62$, p-value $= 0.21$ (0.21 sphericity corrected), $\eta^2 = 0.057$; Block by Sequence interaction: $F(4,108)= 1.45$, p-value $= 0.22$(0.22 sphericity corrected), $\eta^2 = 0.051$). ).

In terms of accuracy, results showed a similar pattern with a main effect of Sequence ($F(2,54) = 6.59$, p-value = 0.0027 (0. 0027 sphericity corrected), $\eta^2 = 0.20$; Figure S6b), yet no effect of Block ($F(2,54) = 0.81$, p-value = 0.45 (0.44 sphericity corrected), $\eta^2 = 0.029$) nor interaction of Block by Sequence were observed ($F(4,108)= 0.24$, p-value = 0.92 (0.89 sphericity corrected), $\eta^2 = 0.0088$).

Collectively, these results suggest that one sequence was differently performed than the other two but as sequences / conditions combinations were pseudo-randomized, it is unlikely that this affected the results presented in the manuscript (and see below for negative control analyses on conditions).

*Equivalent baseline performance between the three conditions*

Similar analyses testing for baseline differences between the reactivated-up, the reactivated-down, and the non-reactivated sequences were performed.

Analyses of the pre-night training data indicated a significant effect of block, but no main effect of condition or block by condition interaction suggesting that participants learned the three sequence conditions to a similar extent during initial learning (21 blocks of training; main effect of Block: $F(20,540) = 42.5$, p-value < 0.001, $\eta^2 = 0.61$; main effect of Condition: $F(2,54)= 0.53$; p-value = 0.59, $\eta^2 = 0.02$; Block by Condition interaction: $F(40,1080)= 1.14$, p-value = 0.26, $\eta^2 = 0.04$; Figure 2). Similar results were observed for accuracy, i.e., a significant main effect of Block ($F(20,540) = 3.0$, p-value < 0.001, $\eta^2 = 0.10$) but no main effect of Condition ($F(2,54) = 0.087$, p-value = 0.92, $\eta^2 = 0.0032$) nor Block by Condition interaction ($F(40,1080)= 1.05$, p-value = 0.38, $\eta^2 = 0.037$, Figure S5a).

Analyses of the pre-night test data showed no main effect of block, condition or block by condition interaction (main effect of Block: $F(2,54) = 1.62$; p-value = 0.21, $\eta^2 = 0.057$; main effect of Condition: $F(2,54) = 0.23$; p-value = 0.80, $\eta^2 = 0.0084$; Block by Condition interaction: $F(3,69)= 1.21$; p-value = 0.31, $\eta^2 = 0.0063$; Figure 2) suggesting that asymptotic performance was reached for all three conditions in the test session that followed initial training run. Similar results were observed for performance accuracy (main effect of Block: $F(2,54) = 0.81$; p-value = 0.45, $\eta^2 = 0.029$; main effect of Condition: $F(2,54) = 0.34$; p-value = 0.69, $\eta^2 = 0.99$; Block by Condition interaction: $F(3,69)= 0.20$; p-value = 0.94, $\eta^2 = 0.0075$; Figure S5a).

*Motor execution performance*

We computed the *overall performance change* for both the sequential SRTT (first 4 blocks of the pre-night training vs. 4 last blocks of post-night training collapsed across sequences) and the pseudo-random version of the SRTT (4 blocks pre-night session vs. 4 blocks post-night session).

Two-tailed paired Student t-test revealed that overall performance changes in performance were significantly higher for the sequential SRTT as compared to the random SRTT (t = -10.2, df = 27, p-value < 0.001; Cohen's d = 1.92). Thus, the RT decrease reported on the sequential SRTT in the result section reflect motor sequence learning rather than a mere improvement in motor execution.

PHASES OF AUDITORY STIMULATION

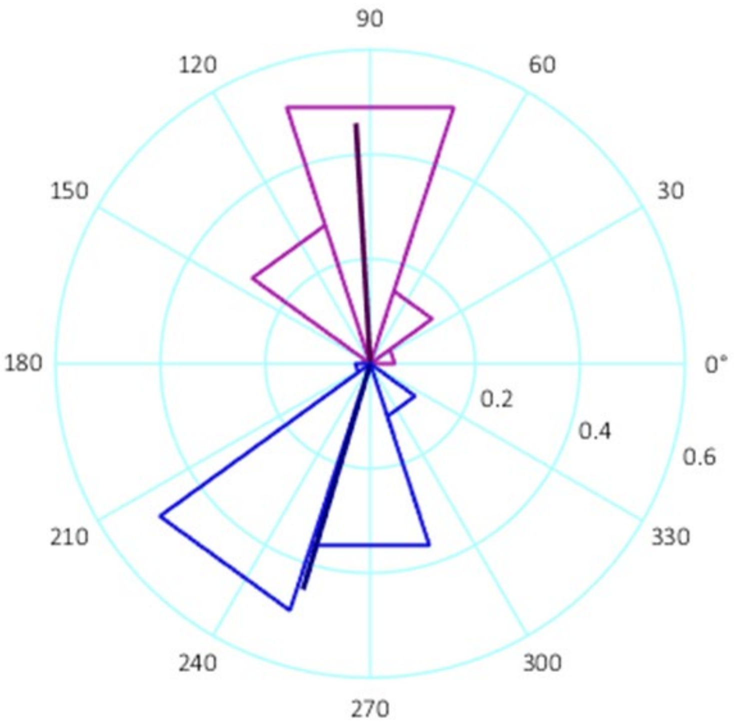

*Figure S7: Stimulation phase. a.* *Density plot of phases in degrees (and mean direction and resultant vector length in bold) of the phase at which up (magenta) and down (blue) auditory cues where sent in the case of true positive trials (% of trials for up and % of trials for down).*

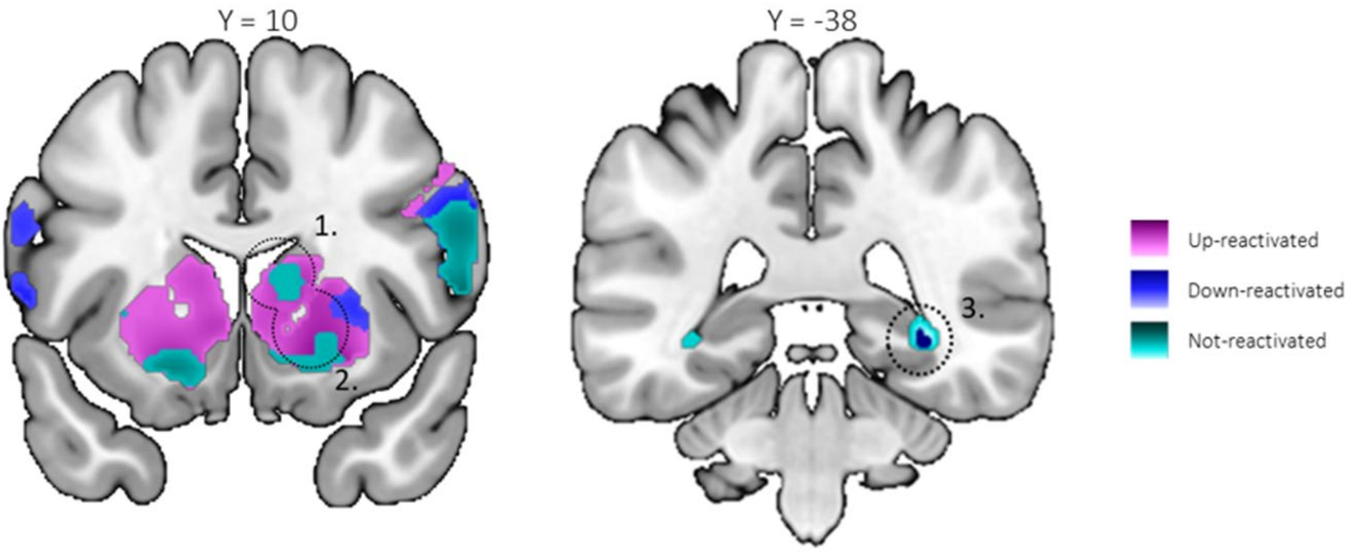

*Figure S8: Seed regions for the connectivity analyses.* Task-related functional connectivity was examined using psychophysiological interaction analyses. Connectivity of three seed regions revealed by the activation-based analyses was assessed. ***a. Striatal seeds (left panel).*** (1) right caudate (x = 10, y = 14, z = 12) and (2) right putamen (x = 18, y = 12, z = -2). ***b. Hippocampal seed (right panel)*** (3) right hippocampus (x = 32, y = -38, z = -6). For each individual, the first eigenvariate of the signal was extracted using Singular Value Decomposition of the time series across the voxels included in a 10 mm-radius sphere centered on these coordinates and shown with the dotted circles. Seed spheres are displayed on a T1-weighted template image super-imposed on the activation maps of the within condition overnight activity increases and decreases with a threshold of $p < 0.005$ uncorrected.

**Table S6**: Coordinates of areas of interest used for spherical small volume corrections for the results presented in the main text.

| Area | x mm | y mm | z mm | Reference |
|---|---|---|---|---|
| HIPPOCAMPUS | | | | |
| | 26 | -34 | -6 | Albouy et al., 2008[12] |
| | 30 | -24 | -15 | Schendan et al., 2003[13] |
| | 24 | -33 | -9 | Schendan et al., 2003[13] |
| | -18 | -16 | -14 | Strange et al., 1999[14] |
| | 24 | -34 | -2 | Strange et al., 1999[14] |
| | 38 | -34 | -2 | Albouy et al., 2008[12] |
| | 34 | -38 | -8 | Albouy et al., 2008[12] |
| | -18 | -34 | 4 | Albouy et al., 2008[12] |
| STRIATUM | | | | |
| Putamen | ±28 | -14 | -8 | Albouy et al., 2008[12] |
| | 29 | -6 | 5 | Lehericy et al., 2005[15] |
| | 27 | 6 | -6 | Schendan et al., 2003[13] |
| | 24 | 10 | -8 | Van Der Graaf et al., 2004[16] |
| Caudate | 20 | -2 | 26 | Albouy et al., 2008[12] |
| | 18 | 8 | 20 | Dolfen et al., 2021[17] |
| | 10 | 26 | 4 | Gann et al., 2021[18] |
| | 15 | 12 | 12 | Schendan et al., 2003[13] |
| | 18 | 22 | 12 | Albouy et al., 2008[12] |
| MOTOR REGIONS | | | | |
| aSPL | ±46 | -38 | 38 | Gann et al., 2021[18] |
| | ±46 | -46 | 46 | Penhune & Doyon 2005[19] |
| M1 | [33 – 39] | [18 – 27] | [51 – 63] | Lehericy et al., 2006[20] |
| | [21 – 40] | [-6 – 11] | [44 – 51] | Paus, 1996[21] |
| | 50 | -18 | 42 | Penhune & Doyon 2005[19] |
| PMC | 50 | -12 | 46 | Penhune & Doyon 2005[19] |
